# Endogenous piRNA-guided slicing triggers responder and trailer piRNA production from viral RNA in *Aedes aegypti* mosquitoes

**DOI:** 10.1101/2020.07.08.193029

**Authors:** Joep Joosten, Gijs J. Overheul, Ronald P. Van Rij, Pascal Miesen

## Abstract

In the germline of animals, PIWI interacting (pi)RNAs protect the genome against the detrimental effects of transposon mobilization. In *Drosophila,* piRNA-mediated cleavage of transposon RNA triggers the production of responder piRNAs via ping-pong amplification. Responder piRNA 3’ end formation is coupled to the production of downstream trailer piRNAs mediated by the nuclease Zucchini, expanding the repertoire of transposon piRNA sequences. In *Aedes aegypti* mosquitoes, piRNAs are generated from viral RNA, yet, it is unknown how viral piRNA 3’ ends are formed and whether viral RNA cleavage gives rise to trailer piRNA production. Here we report that in *Ae. aegypti*, virus- and transposon-derived piRNAs have sharp 3’ ends, and are biased for downstream uridine residues, features reminiscent of Zucchini cleavage of precursor piRNAs in *Drosophila*. We designed a reporter system to study viral piRNA 3’ end formation and found that targeting viral RNA by abundant endogenous piRNAs triggers the production of responder and trailer piRNAs. Using this reporter, we identified the *Ae. aegypti* orthologs of Zucchini and Nibbler, two nucleases involved in piRNA 3’ end formation. Our results furthermore suggest that autonomous piRNA production from viral RNA can be triggered and expanded by an initial cleavage event guided by genome-encoded piRNAs.

## INTRODUCTION

Blood-feeding mosquitoes of the *Aedes* genus are responsible for the transmission of arthropod-borne (arbo)viruses that cause severe diseases, such as dengue, Zika, chikungunya and yellow fever. For efficient transmission to occur, arboviruses have to actively replicate in several mosquito tissues to eventually infect the salivary gland (1). Therefore, suppression of virus replication by the mosquito antiviral immune response strongly affects the efficiency of arboviral spread. The cornerstone of antiviral immunity in insects is the small interfering (si)RNA pathway, in which viral double stranded (ds)RNA is cleaved by Dicer-2 (2). The resulting cleavage products are processed into siRNAs, which provide sequence specificity to the endonuclease Argonaute 2 to direct the cleavage of single stranded viral transcripts. Intriguingly, in *Aedes* mosquitoes, viral RNA is also processed by a somatically active PIWI interacting (pi)RNA pathway, suggesting that two independent small RNA pathways act in parallel to combat viral infections (3).

The piRNA biogenesis machinery has been thoroughly characterized in the model organism *Drosophila melanogaster*, where a gonad-restricted piRNA pathway defends the germline genome from parasitic genetic elements called transposons (4, 5). piRNA biogenesis is initiated by the cleavage of genome-encoded piRNA precursors, which are rich in transposon remnants. Processing of these precursor transcripts into pre-piRNAs is mediated either by a piRNA-guided PIWI protein or the endonuclease Zucchini (Zuc), which operates independently of small RNAs (6–8). Pre-piRNAs are loaded into the PIWI proteins Aubergine (Aub) and Piwi, where their 3’ ends may be further trimmed by the exonuclease Nibbler (Nbr), followed by 2’-*O*-methylation by Hen1 to generate mature piRNAs (9–13). Whereas Piwi translocates to the nucleus to silence transposons at the transcriptional level (14, 15), Aub remains in the cytoplasm where it cleaves (slices) transposon mRNA with sequence complementarity to its associated piRNA (16, 17). The resulting cleavage fragments are loaded into the PIWI protein Argonaute 3 (Ago3) and matured into responder piRNAs by Zuc cleavage and/or Nbr-mediated trimming and subsequent 2′-*O*-methylation by Hen1 (11, 13). In turn, these responder piRNAs direct Ago3-mediated cleavage of piRNA precursors, triggering the production of new initiator piRNAs, completing the so-called ping-pong loop (16–19).

In *Drosophila,* Zuc-mediated generation of piRNA 3’ ends releases a downstream cleavage product that is preferentially loaded into Piwi, thereby generating a new pre-piRNA. This mechanism results in phased processing of piRNA precursor transcripts into a string of piRNAs named trailer piRNAs (7, 8). Thus, the ping-pong loop amplifies those piRNAs that initially recognized active transposons, while phased trailer piRNA production expands the piRNA sequence repertoire for more efficient repression of transposons. Historically, piRNAs derived from cluster transcripts and transposon mRNAs were termed primary and secondary piRNAs, respectively. Hereafter, we use the terms initiator and responder for ping-pong amplified piRNAs and trailer for piRNAs produced through phased biogenesis, as proposed in (4).

The *Aedes aegypti* piRNA pathway is also involved in transposon control and has recently been shown to generate trailer piRNAs (6). However, the pathway differs from that in *Drosophila* in four important ways: *i)* the pathway is active in somatic tissues as well as germline tissues (20, 21), *ii*) the PIWI protein family has expanded to seven members compared to three in *Drosophila* (20,22,23), of which the PIWI proteins Piwi5 and Ago3 engage in ping-pong amplification of piRNAs (24, 25), *iii)* the *Aedes* piRNA pathway processes non-canonical substrates such as viral RNA (24,26,27), and *iv)* mosquito piRNA clusters contain large numbers of endogenous viral elements (EVEs), sequences of non-retroviral RNA viruses inserted in host genomes (28–30). As a consequence, EVEs give rise to abundant piRNAs (28,31–33) and mediate antiviral defense (34, 35). It has been shown that EVE-derived piRNAs can trigger the production of piRNAs from viral RNAs (34, 35). Bases on this observation, we propose that, through trailer piRNA production, a single endogenous initiator piRNA can induce the production of an expanded pool of viral piRNAs, thereby enforcing autonomous piRNA production from viral RNA.

Hitherto, piRNA 3’ end formation and generation of trailer piRNAs have not been studied mechanistically in mosquitoes. Here, we demonstrate that *Ae. aegypti* piRNAs, both of transposon and viral origin, display sequence features indicative of a Zucchini-like biogenesis mechanism. We establish a viral piRNA reporter system to show that AAEL011385 and AAEL005527, the *Ae. aegypti* orthologs of *Drosophila* Zuc and Nibbler, respectively, cooperatively determine piRNA 3’ ends. Furthermore, we demonstrate that cleavage guided by a genome-encoded initiator piRNA triggers the production of trailer piRNAs from the viral genome. We propose that piRNA biogenesis triggered from endogenous sequences, in particular EVEs, may equip *Aedes* mosquitoes with an heritable immune response that, through phasing, is able to adapt to newly encountered and continuously mutating viruses.

## MATERIALS AND METHODS

### Cell culture, dsRNA transfection and infection of Aag2 and U4.4 cells

*Ae. aegypti* Aag2 and *Ae. albopictus* U4.4 cells were maintained in supplemented Leibovitz’s L-15 medium (Invitrogen) at 25°C. For knockdown experiments, dsRNA was transfected using X-tremeGENE HP DNA Transfection Reagent (Roche) according to the manufacturer’s instructions. Where indicated, cells were infected with Sindbis virus (SINV) at a multiplicity of infection (MOI) of 0.1. For further details, see Supplemental Information.

### Generation of reporter viruses

Target sites for *gypsy*- and EVE-initiator piRNAs and trailer cassettes were introduced into an infectious cDNA clone of Sindbis virus downstream of a duplicated subgenomic promoter. Site-directed mutagenesis was used to introduce target site mutations. Subsequently, viruses were grown as described previously (27). For details, see Supplemental Information.

### RNA isolation, RT-qPCR and small RNA northern blotting

Total RNA was isolated using RNA-SOLV reagent (Omega Bio-tek). For RT-qPCR analyses, RNA was DNaseI treated, reverse transcribed, and PCR amplified in the presence of SYBR green. For small RNA northern blotting, RNA was resolved by denaturing urea polyacrylamide gel electrophoresis and transferred to nylon membranes. See Supplemental Information for experimental details and oligonucleotide sequences.

### Generation of small RNA deep sequencing libraries and bioinformatic analyses

For the analyses of small RNAs, deep sequencing libraries were generated using the NEBNext Small RNA Library Prep Set for Illumina (E7560, New England Biolabs) and sequenced on an Illumina Hiseq4000. Sequence data have been deposited in the NCBI sequence read archive under SRA accession SRP272125. Sequencing data were analyzed in Galaxy (36). Reads were mapped to Sindbis virus genomes, transposon sequences, *Ae. aegypti* transcripts, the Phasi Charoen like virus genome and pre-miRNA sequences using Bowtie (37). Further details are provided in the Supplemental Information.

### Immunofluorescence analyses of Zuc localization

Aag2 cells were transfected with a plasmid expressing 3×flag tagged Zuc using X-tremeGENE HP DNA Transfection Reagent (Roche), and fixed 48 hours after transfection. Cells were incubated with a mouse anti-flag antibody (Sigma, F1804, RRID: AB_262044), followed by goat anti-mouse IgG Alexa fluor 568 (Invitrogen, A-11004, RRID: AB_2534072). Mitochondria were stained using Mitoview Green (Biotium). For further information, see Supplemental Information.

### Immunoprecipitation and western blot

A 3×flag tagged Zuc expression plasmid was transfected into Aag2 cells using X-tremeGENE HP DNA Transfection Reagent (Roche). Cells were lysed and lysates incubated with M2-Flag beads (Sigma) to immunoprecipitate 3×flag tagged Zuc and interacting proteins. For western blot analyses, samples were resolved on polyacrylamide gels, blotted to nitrocellulose membranes and stained with the following antibodies generated in our laboratory (25, 38): rabbit-anti-Ago3, -Piwi4, -Piwi5 and - Piwi6 (all at 1:500), and mouse anti-flag (1:1000, Sigma, F1804, RRID: AB_262044). Subsequently, goat-anti-rabbit-IRdye800 [Li-cor; 926-32211, RRID: AB_621843] and goat-anti-mouse-IRdye680 [926-68070, RRID: AB_10956588] were used for visualization. Small RNAs were isolated from PIWI protein immunoprecipitates as described in (25). For experimental details, see Supplemental Information.

### Statistical analyses

Unless indicated otherwise, unpaired two tailed t-tests with Holm-Sidak correction for multiple comparisons were used for statistical analyses (* *P* < 0.05; ** *P* < 0.005; *** *P* < 0.0005) using Prism 8 (GraphPad Software). For statistical analysis of sharpness scores in Figures 4E-F, see Supplemental Information.

## RESULTS AND DISCUSSION

### *Aedes aegypti* piRNAs have sharp 3’ ends

In *Drosophila*, piRNA 3’ end formation is largely dependent on the cleavage of pre-piRNAs by the endonuclease Zucchini (Zuc). Zuc uses a sequence motif to preferentially cleave upstream of uridine residues *in vivo* (39–41), hence, piRNAs generated by Zuc generally have sharp 3’ ends and the nucleotide directly downstream of the 3’ end is biased towards uridine (+1U bias) (7, 8). We examined whether these characteristics were present in our previously generated small RNA deep sequencing libraries from *Ae. aegypti* Aag2 cells infected with Sindbis virus (SINV) (24). We first analyzed transposon-derived piRNAs and found that piRNAs that shared the same 5’ end generally had the same length (Figure 1A). Specifically, for almost 60% of piRNAs the dominant length made up more than 75% of sequenced reads. We selected these piRNAs and inspected the identity of the nucleotides downstream of that most abundant piRNA isoform. We found that the nucleotide position directly following the 3’ end of the piRNA was biased for uridine (Figure 1B), strongly indicating that these piRNAs were generated by a mechanism resembling Zuc cleavage in *Drosophila*. We next analyzed the characteristics of 3’ ends of viral (v)piRNAs derived from the SINV genome. SINV is a positive-strand RNA virus of the *Togaviridae* family. During its replication cycle, genomic sense (+) strand RNA serves as a template for the production of antigenomic antisense (-) strand RNA, which in turn provides a template for production of genomic and subgenomic RNA species (42). Strikingly, sharp 3’ ends were clearly visible for vpiRNAs, irrespective of the strand from which the piRNAs were produced (Figure 1C). In addition, a clear +1U bias was observed, especially for antisense strand derived piRNAs (Figure 1D). These findings suggest that 3’ ends of both Ago3-associated (+) strand derived vpiRNAs and Piwi5-bound (-) strand derived vpiRNAs (24, 25), are generated, at least in part, by Zuc-like cleavage events. Interestingly, we also observed sharp 3’ ends and +1U biases for vpiRNAs generated from Phasi Charoen-like virus (Figure S1A-B), a negative-strand RNA virus from the *Phenuiviridae* family that persistently infects Aag2 cells (43). These findings indicate that a Zuc- like biogenesis mechanism contributes to 3’ end formation of piRNAs derived from transposons, as well as diverse RNA viruses.

**Figure 1.**
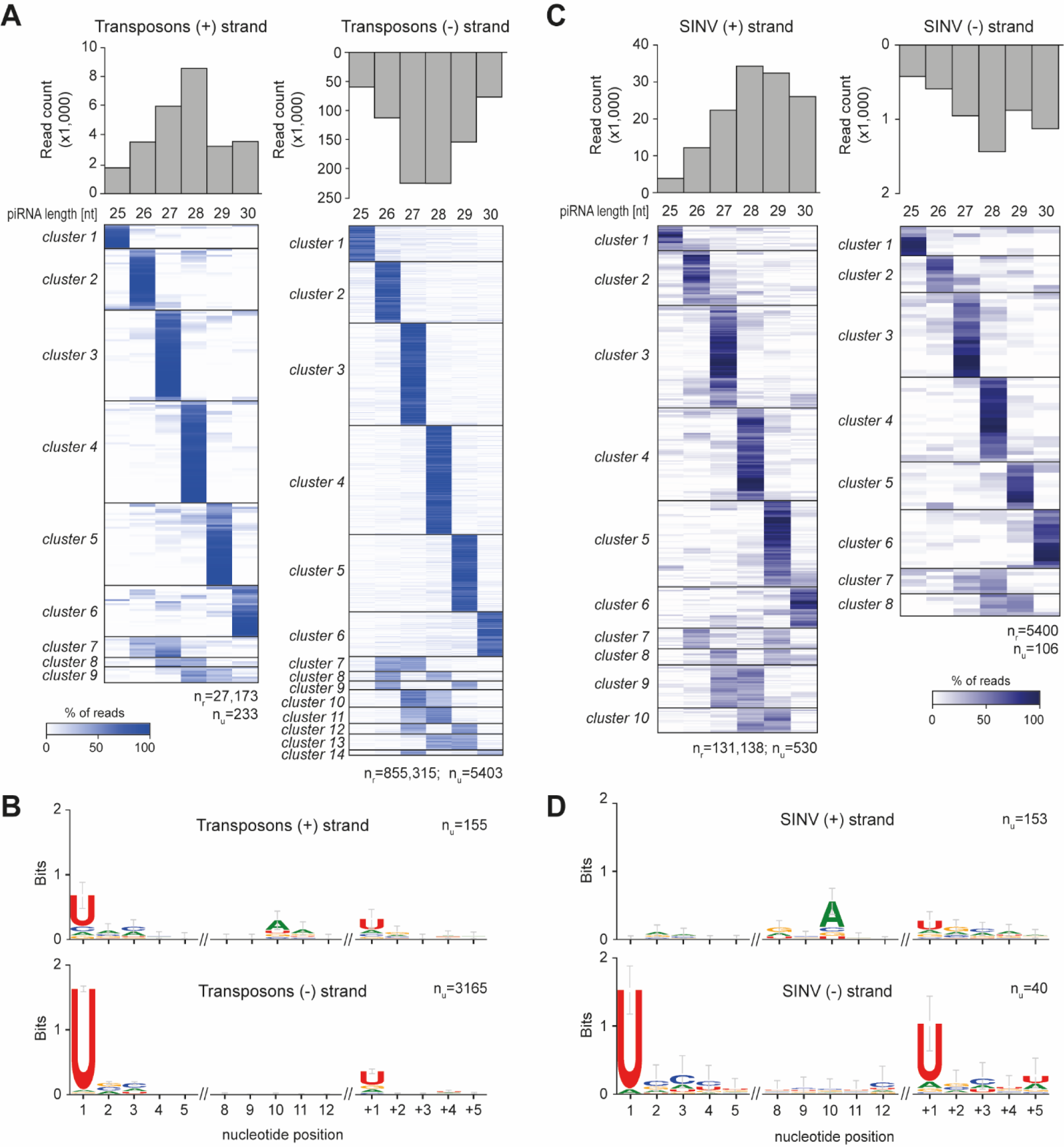
*Aedes aegypti* piRNAs have sharp 3’ ends. **(A)** Analysis of individual piRNA lengths. The bar graphs show the size distribution of transposon-derived sense ((+), left) and antisense ((-), right) piRNAs from the combined three dsLuc small RNA libraries published in (24). The heat maps show the relative size distribution of transposon-derived piRNAs that share the same 5’ end (one 5’ end is one line in the heat map). Shades of blue indicate the percentage of reads contributing to the indicated piRNA length, white represents absence of reads of the specific size. The number of reads (nr) and the number of unique piRNAs 5’ ends (nu) that underlie the heat map are indicated. A minimum of 20 reads per unique piRNA position was required to be included in the analysis. **(B)** Nucleotide biases at the indicated positions of transposon-derived piRNAs and the sequence at the genomic region directly downstream (+1 until +5) of the piRNA 3’ ends. Only piRNAs from (A) that had a dominant piRNA length (at least 75% of reads were of the same size) were considered in this analysis and only unique piRNA sequences were analyzed, irrespective of read count. nu indicates the number of sequences underlying the sequence logo. **(C-D)** The same analysis as for A and B was applied to SINV-derived piRNAs.

### Genome-encoded piRNAs trigger production of virus-derived responder piRNAs

To study vpiRNA 3’ end formation, we designed a SINV-based reporter system which contained a duplicated subgenomic promoter driving the expression of a non-coding RNA sequence that harbors a target site for an abundant initiator piRNA (referred to as reporter cassette, Figure 2A and S2A). These Piwi 5-associated initiator piRNAs (Figure S2B-C) either derived from the Ty3/*gypsy* LTR retrotransposon *gypsy73* (Figure 2A) or from an EVE sequence of flaviviral origin (Figure S2A; see also Supplemental text). From here on, we will refer to these piRNAs as *gypsy* and EVE initiator piRNAs. Initiator piRNA-guided recognition of the artificial target site in the reporter virus is expected to trigger slicing by Piwi5 and subsequent processing of the resulting cleavage fragment into an Ago3- associated responder piRNA through ping-pong amplification. Indeed, virus-derived responder piRNAs were abundantly produced in Aag2 cells infected with the reporter viruses containing the artificial piRNA target sites but not in uninfected cells and cells infected with a control virus expressing GFP from the duplicated subgenomic promoter (SINV 3’ GFP) (Figure 2B; S2D). These results indicate that endogenous piRNAs can instruct the cleavage of exogenous viral RNA and induce the production of responder piRNAs during acute infection.

**Figure 2.**
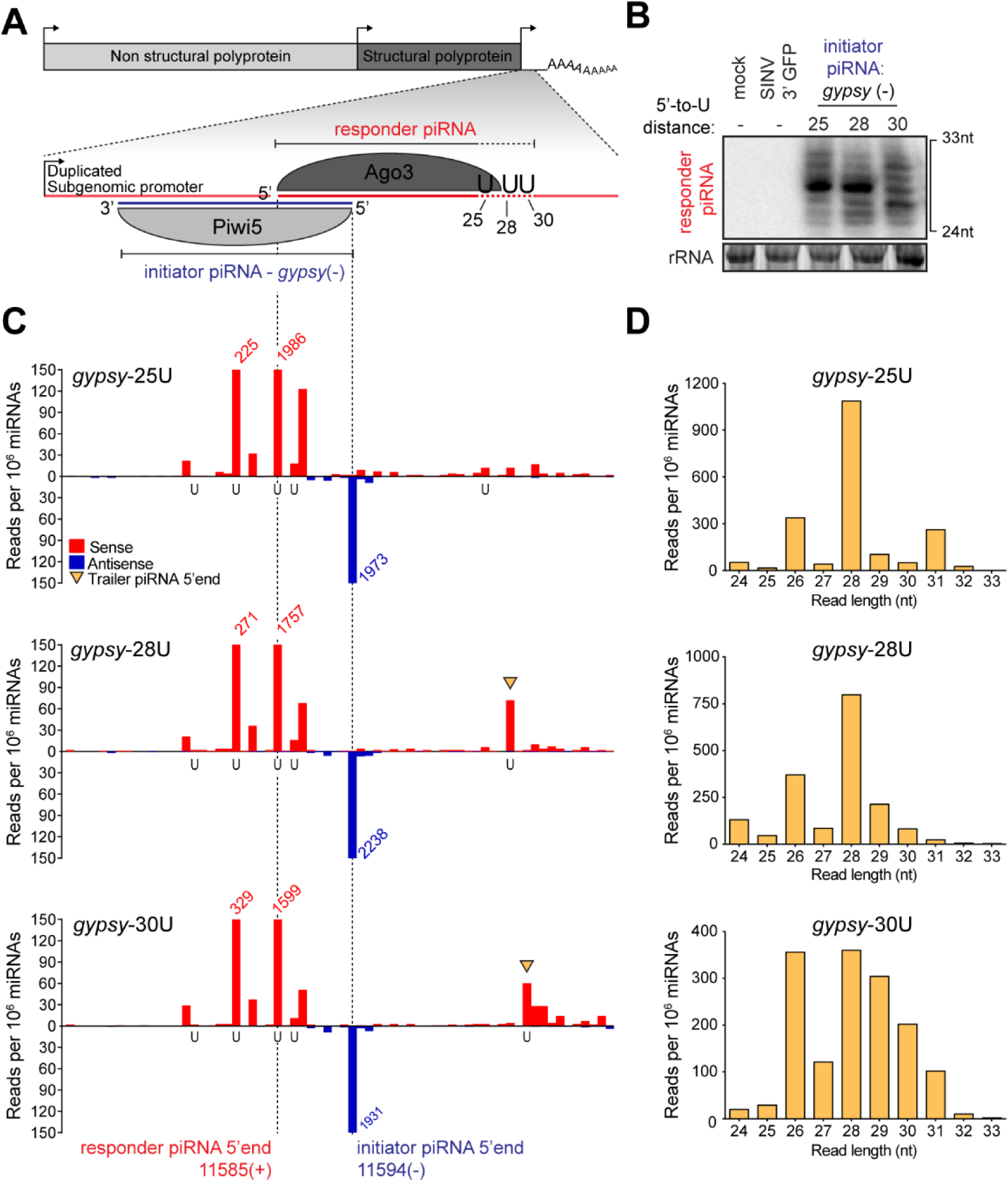
An endogenous piRNA is able to trigger production of virus-derived responder piRNAs. **(A)** Schematic representation of recombinant Sindbis reporter viruses. The enlarged view depicts the reporter locus expressed under the control of a duplicated subgenomic promoter. This non-coding RNA harbors a target site for a Piwi5-associated initiator piRNA derived from the *Ty3*-*gypsy* retrotransposon (*gypsy* - indicated in blue). Slicing of this target triggers the production of responder piRNAs (indicated in red) that are loaded into Ago3. The positions of downstream uridine residues in the various viruses used in later experiments are shown. **(B)** Northern blot analyses of responder piRNAs in Aag2 cells infected with the indicated reporter viruses. Numbers indicate the distance between the 5’ end of the responder piRNA and the first downstream uridine residue. SINV 3’ GFP is a virus without an initiator piRNA target site and serves as a negative control, as do mock infected cells. The positions marking 24 and 33 nt are inferred from an EtBr- stained small RNA size marker. EtBr-stained rRNA serves as loading control. The northern blot is representative of four independent experiments. **(C)** Visualization of 5’ ends of sense (red) and antisense (blue) piRNAs (24-33 nt) mapping to the reporter locus of the indicated viruses. Dashed lines indicate initiator and responder piRNA 5’ ends and the positions of grey shading indicate the location of uridine residues on the sense strand of the various viruses. The red and blue numbers show piRNA counts (in reads per 10^6^ miRNAs) that exceed the range of the y-axis; yellow arrowheads indicate the 5’ ends of trailer piRNAs. For each virus infection, a single sequencing library was analyzed (same for (D)). **(D)** Size distribution of responder piRNAs (SINV genome position 11585 (+)) produced from the various *gypsy*-viruses, as determined by small RNA deep sequencing. Read counts were normalized to the number of miRNAs in each library.

Our previous results indicated that both transposon- and SINV-derived piRNAs have a strong bias towards a uridine residue directly downstream of their 3’ ends (Figure 1B, D). To study the importance of the +1U position for viral responder piRNA production, we introduced uridine residues at specified distances from the putative Piwi5 slice site in the viral reporter (Figure 2A; viruses were named *gypsy*- 25U, 28U, and 30U according to the distance of responder piRNA 5’ end to the +1U). Responder piRNAs were readily detected by high resolution northern blotting for all reporter viruses (Figure 2B), yet the size of the responder piRNA did not reflect the distance between the Piwi5 cleavage site and the downstream uridine residue. While no clear differences in responder piRNA size distribution were observed between the *gypsy*-25U and *gypsy*-28U viruses, increasing the 5’ end-to-U distance to 30 nt (*gypsy*-30U) resulted in a more diffuse pattern of responder piRNA lengths (Figure 2B). These data suggest that downstream uridines may not be the only determinant for 3’ end formation of the reporter- derived responder piRNAs or that additional exonucleolytic trimming of pre-piRNA 3’ ends masked a putative endonucleolytic cleavage event directly upstream of the uridine residues. To discriminate between these two possibilities, we analyzed small RNA sequences from Aag2 cells infected with the various reporter viruses.

Mapping of piRNA 5’ ends to the genomes of *gypsy*-targeted viruses revealed that virtually no antisense piRNAs other than the initiator piRNAs map to the reporter sequence (Figure 2C). These initiator piRNAs triggered the production of highly abundant sense responder piRNAs with a characteristic 10 nt overlap of piRNA 5’ ends, indicative of production by ping-pong amplification (Figure 2C). Responder piRNA size distribution for *gypsy*-targeted viruses generally recapitulated the results from the northern blot analysis, with a broader size distribution for the *gypsy*-30U virus (Figure 2D).

In viruses with a distance of responder piRNA 5’ ends to +1U ≥28 nt, we detected putative trailer piRNAs downstream of the responder piRNA (indicated with yellow arrowheads in Figure 2C). Strikingly, the 5’ end of these trailer piRNAs was sharply defined by the position of the downstream uridine. Similarly, we observed U-directed trailer piRNA production for the EVE-triggered viruses (Figure S2E, Supplemental text). These data suggest that downstream uridines may instruct the positioning of endonucleolytic cleavage, thus coupling responder piRNA 3’ end formation to trailer piRNA 5’ end formation, as previously described in *Drosophila* (7, 8). The heterogeneous responder piRNA size likely results from subsequent exonucleolytic trimming.

Intriguingly, only minor U-directed trailer piRNA production was observed in cells infected with the *gypsy*-25U virus. We speculated that the uridine at position 25 may be covered by the Ago3 protein, rendering it inaccessible for endonucleolytic cleavage (Figure 2A). In line with this hypothesis, we observed that only very few responder piRNAs <26 nt are produced from any *gypsy* triggered reporter virus (Figure 2D). Furthermore, small RNA sequencing data from Ago3 immunoprecipitates (IP) indicated a clear preference of Ago3, Piwi5 and Piwi6 to bind piRNAs in the size range of 26-30 nt (Figure S3A,C,D), whereas the Piwi4 IP library was dominated by tapiR1, a highly abundant piRNA that is 30 nt in size (38) (Figure S3B). These data strongly support the notion that the lack of trailer piRNAs in the *gypsy*-25U reporter virus is explained by inaccessibility of the introduced U residue due to steric hindrance by the associated PIWI protein. Altogether, these data indicate that our viral reporter system faithfully recapitulates various aspects of piRNA 3’ end formation, serving as an amenable tool to study responder and trailer piRNA biogenesis.

### Responder piRNAs are produced through ping-pong mediated slicing

We previously identified Ago3 and Piwi5 as the core components of the ping-pong amplification loop in *Ae. aegypti* (24, 25). We therefore set out to validate that these PIWI proteins are responsible for the generation of the responder piRNAs from our reporter viruses. First, we determined the levels of *gypsy* and EVE initiator piRNAs in previously published small RNA deep sequencing libraries generated from Aag2 cells in which somatic PIWI proteins (Ago3 and Piwi4-6) were depleted (24). In accordance with ping-pong dependent production, the level of the *gypsy* initiator piRNA was significantly reduced upon knockdown of Ago3 and Piwi5 (2.2- and 5.5-fold, respectively; Figure 3A). Similarly, EVE-derived initiator piRNA levels were significantly reduced upon knockdown of the ping-pong partners Ago3 and Piwi5 (1.6-fold and 2.3-fold, respectively, Figure 3B). Unexpectedly, while Piwi4 depletion had previously been reported to cause a decline of piRNAs from a large proportion of transposons (24, 34), knockdown of Piwi4 resulted in an almost twofold increase in *gypsy* initiator piRNAs (Figure 3A). Moreover, Piwi4 knockdown caused a general increase in piRNA expression from the entire genomic locus that produces the *gypsy* initiator piRNA (Figure S4A). This intriguing finding suggests that Piwi4 controls the expression of selected piRNA cluster transcripts, the mechanism of which requires further investigation.

**Figure 3.**
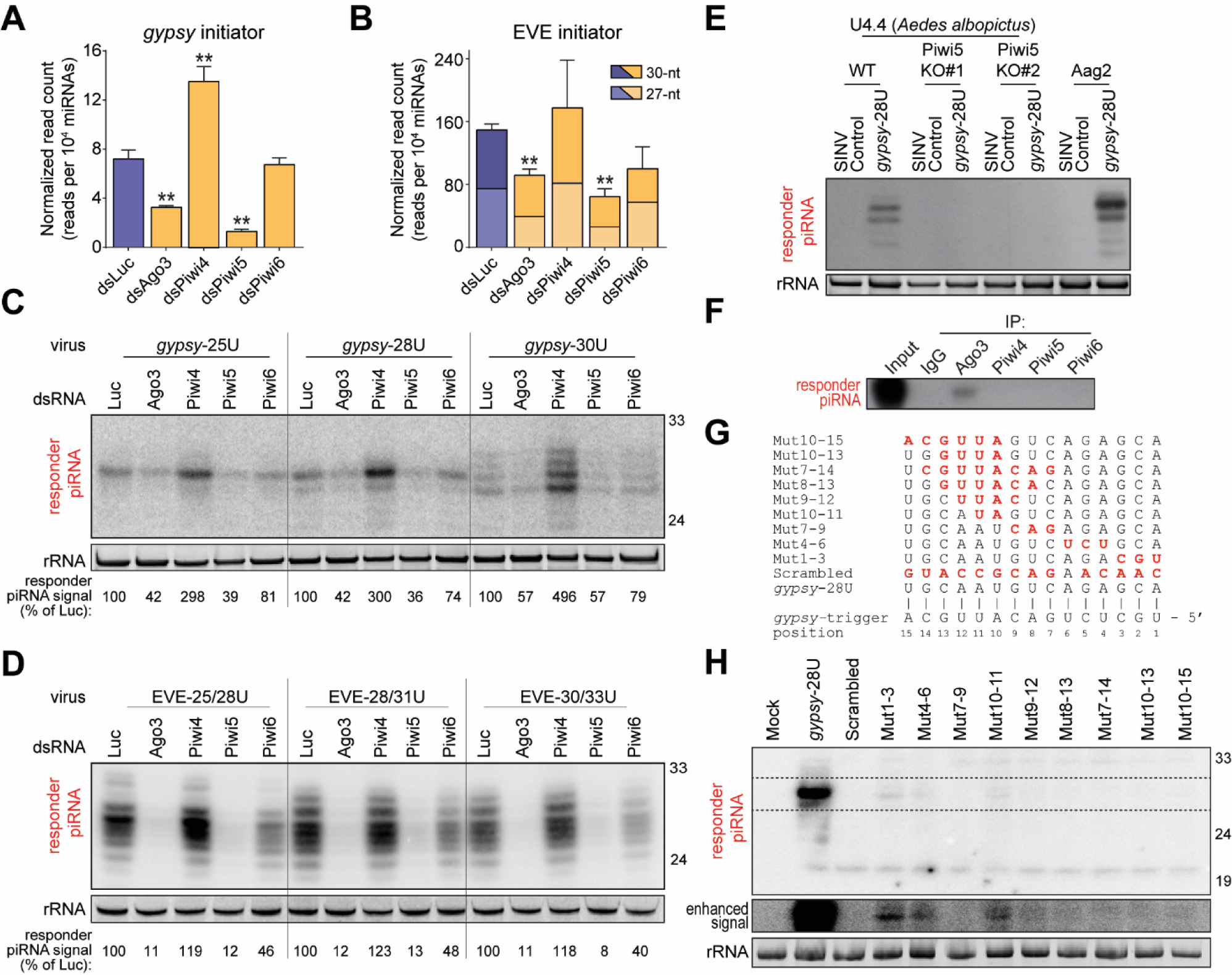
Responder vpiRNAs are produced through ping-pong mediated slicing. **(A-B)** Levels of the *gypsy*- (A) and EVE-derived (B) initiator piRNAs in previously published small RNA sequencing libraries generated from Aag2 cells in which indicated PIWI proteins were depleted (24). The flavivirus EVE-derived piRNA exists as 27-nt and 30-nt isoforms; the error bars and statistical analyses are based on combined counts of the two isoforms. Asterisks denote statistical significance as determined by unpaired two tailed t-tests with Holm-Sidak correction (** *P* < 0.005). Bars and whiskers represent the mean and SD of three independent libraries, respectively. **(C-D)** Northern blot analysis of viral responder piRNAs in Aag2 cells in which indicated PIWI proteins were depleted. Cells are infected with indicated *gypsy* (C) and EVE (D) targeted viruses. EtBr-stained rRNA serves as loading control. Numbers indicate responder piRNA signals, normalized to the loading control, as a percentage of the responder piRNA signal in dsLuc treated cell infected with the same virus. For both (C) and (D), n = 1 biological replicate. **(E)** Northern blot analysis of responder piRNAs in wildtype (WT) and two independent Piwi5 knockout (KO) *Ae. albopictus* U4.4 cell lines, and in *Ae. aegypti* Aag2 cells infected with the *gypsy*-28U virus or a virus that does not express the reporter locus from the second subgenomic promoter (SINV Control). EtBr-stained rRNA serves as a loading control. Northern blot is a representative of two independent biological replicates. **(F)** Northern blot analysis of the viral responder piRNA in PIWI protein immunoprecipitates (IP) from Aag2 cells infected with the *gypsy-* U interval virus (n = 1 biological replicate). The *gypsy*-U interval virus, described in detail in Figure 6, contains the same initiator piRNA target site and responder sequence as the *gypsy-*28U virus. As a control, non-specific rabbit IgG was used for IP. **(G)** Overview of target site mutations for the various viruses shown in (H). Red bold font indicates residues that are mismatched with the gypsy initiator piRNA and the numbers denote positions relative to the *gypsy* initiator piRNA 5’ end. Northern blot analysis of responder piRNAs in Aag2 cells infected with indicated (mutant) viruses (n = 1 biological replicate). The dashed box denotes an area for which the contrast was adjusted to enhance weak responder piRNA signals (enhanced signal – middle panel). The ‘minimal responder’ probe used in this experiment hybridizes to the last 18 nt of the 3’ end of responder piRNAs, which are identical for all viruses (see also Figure S4G). EtBr stained rRNA serves as loading control (bottom panel).

We next assessed the effect of PIWI knockdown on viral responder piRNA levels. As expected, responder piRNA production from the *gypsy*-targeted viruses was reduced upon knockdown of genes encoding the ping-pong partners Ago3 and Piwi5 and, to a lesser extent, Piwi6 (Figure 3C), despite moderate knockdown efficiency (Figure S4B). No significant effects on viral RNA levels were observed for any of these PIWI knockdowns, indicating that reduced piRNA levels were not due to reduced viral replication (Figure S4C). Higher levels of the *gypsy*-derived initiator piRNA (Figure 3A) likely explains the observed increase in responder piRNA production upon Piwi4 knockdown (Figure 3C). We obtained similar results for the reporter viruses that are targeted by the EVE-derived initiator piRNA. Efficient knockdown of the ping-pong partners Ago3 and Piwi5 resulted in a dramatic decline in responder piRNA production from these viruses, while Piwi6 knockdown had a moderate effect (Figure 3D, S4D). No effects of PIWI knockdown on viral RNA replication were observed (Figure S4E). Importantly, Piwi4 knockdown, which did not result in altered EVE initiator piRNA levels (Figure 3B), barely affected responder piRNA production (Figure 3D), suggesting that Piwi4 has no direct involvement in ping-pong amplification of responder piRNAs. This is in line with previous findings that Piwi4 does not associate with Ago3 and Piwi5 (44) and is not required for ping-pong amplification of piRNAs (24,45,46). The Piwi5-dependency of *gypsy* responder piRNA production was further validated in Piwi5 knockout U4.4 cells, derived from the closely related mosquito *Aedes albopictus* (Figure 3E, S4F). Moreover, as expected from their ping-pong dependent production, responder piRNAs were specifically bound to Ago3 (Figure 3F).

We next investigated base-pairing requirements for responder piRNA production by introducing mutations into the seed region (nt 2-8), and around the putative slice site (nt 10-11) of the *gypsy* piRNA target site (Figure 3G, S4G). Responder piRNA production was strongly depleted in viruses in which mutations were introduced in the seed sequence (Mut 1-3, Mut 4-6 and Mut 7-9) compared to a virus bearing the intact target site (*gypsy*-28U, Figure 3H), indicating that seed-based target recognition is required for efficient responder piRNA production. Similarly, introducing mismatches around the slice site (Mut 10-11, Mut 9-12, Mut 10-13, Mut 10-15, Mut 8-13 and Mut 7-14) resulted in strongly reduced responder piRNA production. As viral RNA levels are virtually unchanged between all viruses, reduced responder piRNA production cannot result from differences in the amount of available substrate (Figure S4H). Weak responder piRNA production was observed in two seed mutants (Mut 1-3 and Mut 4-6) and a slice site mutant (Mut10-11), suggesting that low level slicing may occur even in the absence of full complementarity in the seed region or the slice site, in line with earlier findings in *Drosophila* (8), mice (47) and mosquitoes (48). Altogether, these data show that slicing by the ping-pong partners Ago3 and Piwi5 is required for the production of responder piRNAs from the viral reporter.

### Zuc-mediated endonucleolytic cleavage defines piRNA 3’ends

The presence of sharply defined piRNA 3’ ends in combination with a bias for a downstream uridine (Figure 1) suggest that a Zuc-like endonuclease generates responder piRNAs 3’ ends and trailer piRNA 5’ ends. As the nuclease activity of Zuc lies in its phospholipase D (PLDc_2)-domain (39, 40), we aligned the sequences of all *Ae. aegypti* PLDc_2-domain containing proteins with those of Zuc orthologs from fruit flies, silkworm and mouse (*Dm*Zuc, *Bm*Zuc and *Mm*MitoPLD, respectively) and found that AAEL011385 had the highest similarity to the various Zuc orthologs (Figure 4A). The protein encoded by this gene contains a fully conserved catalytic H(X)K(X4)D (HKD)-motif (Figure S5A), suggesting that it is a functional endonuclease. Moreover, akin to Zuc orthologs in various other species (4, 5), AAEL011385/Zuc localized to the mitochondria in Aag2 cells (Figure 4B).

**Figure 4.**
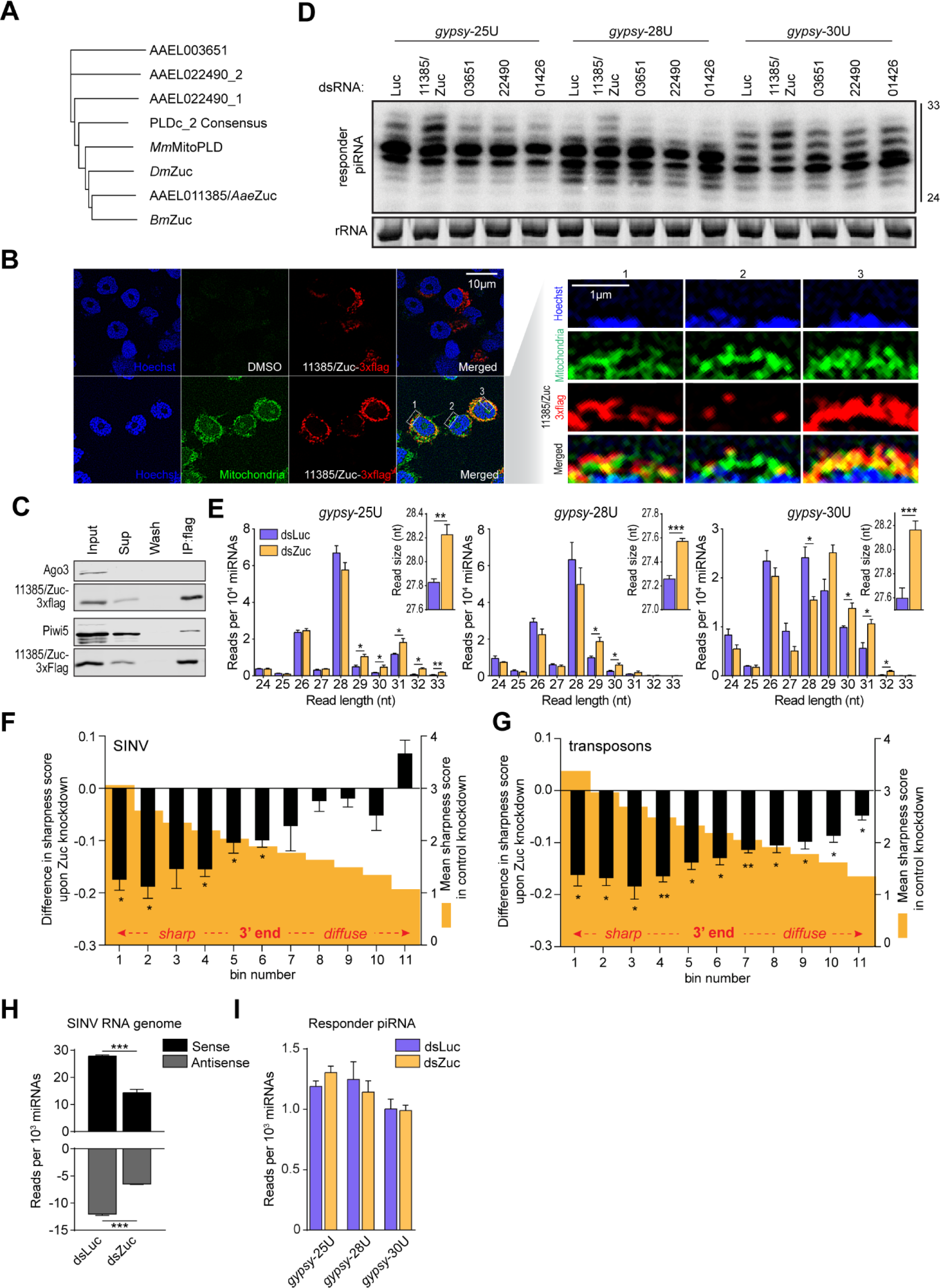
Zuc-mediated endonucleolytic cleavage defines vpiRNA 3’ends. **(A)** Phylogenetic tree based on PLDc_2 domains of *Ae. aegypti* PLDc_2 domain containing proteins and Zuc orthologs from *Drosophila (Dm* Zuc), silkworm (*Bm*Zuc) and mouse (*Mm*MitoPLD). **(B)** Confocal microscopy images of Aag2 cells expressing 3×flag tagged AAEL011385/Zuc. Mitochondria were stained using Mitoview green. Enlargements of the areas indicated by dashed boxes are shown in the right panels, with the nuclei oriented at the bottom. Scale bars denote 10µm (left panel) and 1µm (right panel). **(C)** Western blot of indicated proteins in AAEL011385/Zuc-3×flag immunoprecipitation (IP). **(D)** Northern blot analysis of viral responder piRNAs in Aag2 cells infected with the indicated viruses upon knockdown of *Ae. aegypti* PLDc_2 domain containing proteins and the PARN ortholog AAEL001426. Numbers indicate the VectorBase gene identifiers (without the AAEL0 prefix). The 24 and 33 nt size markers are inferred from an EtBr stained small RNA marker and rRNA stained by EtBr served as a loading control. The knockdown screen has been performed once; the AAEL011385/Zuc knockdown phenotype has been observed in eight independent infections. **(E)** Size distribution of viral responder piRNAs (SINV genome position 11585(+)) in small RNA sequencing libraries from Aag2 cells treated with dsRNA targeting luciferase (dsLuc) and Zuc (dsZuc). Read counts were normalized to the number of miRNAs in each library. The inset shows the average responder piRNA read size in Luc- and Zuc knockdown libraries. Bars and whiskers represent mean and SD of three independent libraries, respectively. Asterisks denote statistical significance as determined by unpaired two tailed t-tests with Holm-Sidak correction (* *P* < 0.05, ** *P* < 0.005, *** *P* < 0.0005). **(F-G)** Distribution of sharpness of piRNA 3’ ends upon Zuc knockdown. A sharpness score was attributed to the 275 most abundant viral piRNAs upstream of the artificial reporter cassette (F) and 4400 most abundant transposon piRNAs (G). The maximum score (3.81) is reached if 100% of piRNA reads that share the same 5’ end have the same length. For each piRNA, the sharpness score was determined in control (Luc) and Zuc knockdown conditions. The piRNAs were ranked and binned (n=25 vpiRNAs and n=400 transposon piRNAs per bin, respectively) according to the score in the control knockdown. For each bin, the mean sharpness score of all nine control knockdown libraries (three for each type of reporter virus) are plotted (orange shade, right y-axis). The difference of piRNA sharpness score upon Zuc knockdown was calculated and averaged for each type of reporter virus separately. Plotted is the mean and SEM of these average scores (left y-axis). A two-sided student’s t-test was applied to each bin to assess whether its mean was significantly different from zero. * *P* < 0.05 and ** *P* < 0.005. **(H)** Read count of piRNAs mapping to the SINV genomic RNA in dsLuc and dsZuc treated Aag2 cells. Average piRNA counts were calculated from three independent libraries per virus infection. As the SINV genomic RNA is common for the three reporter viruses, these averages were used to determine mean piRNA read counts +/- SEM (bars and whiskers, respectively). Statistical significance determined by unpaired two tailed t-tests with Holm-Sidak correction is indicated with asterisks (*** *P* < 0.0005). **(I)** Responder piRNA levels in the reporter locus in dsLuc and dsZuc treated Aag2 cells infected with the indicated viruses. Bars and whiskers show the mean +/- SD of three independent libraries.

In *Drosophila*, a strong interaction between Zuc and Aub (the ortholog of Piwi5), as well as weak associations between Zuc and Ago3/Piwi have been observed (49–51). We thus evaluated whether AAEL011385/Zuc interacts with somatic *Aedes aegypti* PIWI proteins. While we readily detect Piwi5 in AAEL011385/Zuc immunoprecipitates, we did not observe interaction of AAEL011385/Zuc with Ago3 (Figure 4C), nor with Piwi4 and Piwi6 (Figure S5B).

To our surprise, we found that AAEL011385/Zuc contains a sizeable insertion directly downstream of the catalytic HKD-motif (Figure S5A). Moreover, during cloning of the AAEL011385/Zuc gene, we found that the size of this insertion is further increased by an additional 32 amino acids in Aag2 cells (Figure S5A). Relative to mouse mitoPLD, *Drosophila and Bombyx* Zuc also contain (much smaller) insertions at the same position, which corresponds to the location of a helix that sticks out of core structure of *Drosophila* Zuc (40). Multiple sequence alignment revealed that a large insertion of >40 amino acids is present in *Culicidae* (mosquitoes) but not in other vector species such as ticks, lice, and tsetse flies, indicating that it is a variable region beyond *Aedine* mosquitoes, the function of which remains to be understood.

Using our viral responder piRNA reporter, we next aimed to validate AAEL011385 as the functional ortholog of *DmZuc*. Indeed, knockdown of AAEL011385/Zuc in Aag2 cells resulted in longer viral responder piRNAs with a broader size distribution (Figure 4D, Figure S5C), which is consistent with less efficient piRNA 3’ end generation. Knockdown of the other PLDc_2 domain containing proteins (AAEL003651 and -22490) did not affect responder piRNA size (Figure 4D), despite very efficient knockdown (88-94% and 96-98% for AAEL003651 and -22490, respectively, compared to 58-71% for AAEL011385/Zuc; Figure S5D). Viral RNA levels were not consistently affected by knockdown of any of the genes tested (Figure S5E). Small RNA deep-sequencing of dsAAEL011385/Zuc-treated Aag2 cells recapitulated the phenotype seen by northern blotting, with a general increase in piRNA length upon knockdown of AAEL011385/Zuc (Figure 4E). Based on these results, we conclude that AAEL011385 is the functional Zuc ortholog in *Ae. aegypti*.

We next studied the general effect of Zuc knockdown on the 3’ end sharpness on the entire population of vpiRNAs outside of the reporter cassette. Therefore, we calculated sharpness scores for vpiRNAs that share the same 5’ end based on the Shannon entropy of their size distribution. A high score indicates that piRNAs with identical 5’ ends generally also had the same length whereas a lower score indicates a more diffuse size distribution. As expected, Zuc knockdown significantly reduced sharpness scores of vpiRNAs, in particular for those that had the sharpest 3’ ends in the control knockdown and were therefore likely the most dependent on Zuc cleavage (Figure 4F). The same effect was observed for piRNAs that mapped to transposon sequences (Figure 4G). Moreover, Zuc knockdown resulted in an increase in size of piRNAs produced from substrates of various origins, including transposons, mRNAs and viral RNAs (Figure S5F), suggesting that Zuc is ubiquitously active in the formation of piRNA 3’ends and is important for maturation of piRNAs from a broad repertoire of RNA substrates.

We next assessed the effect of Zuc depletion on overall vpiRNA levels. As expected, Zuc depletion reduced overall vpiRNA production from the SINV genomic and subgenomic RNA, which was common to the *gypsy*-25U, -28U and -30U viruses (Figure 4H). Yet, the abundance of the *gypsy*- triggered responder piRNA produced from the artificially introduced reporter locus was not affected by Zuc knockdown (Figure 4I), suggesting that an alternative mechanism contributes to 3’ end formation of this particular piRNA. We propose that upon Zuc knockdown, the Ago3-bound piRNA precursor is cleaved downstream of the uridine residue, either by a hitherto unknown endonuclease or by other PIWI-piRNA ribonucleoprotein complexes, as previously reported in *Drosophila* (11).

### A subset of responder piRNAs undergoes Nibbler-mediated trimming

In *Drosophila*, piRNA 3’ ends are generated by the concerted activities of two enzymes: the endonuclease Zuc (6–8) and the 3’ – 5’ exonuclease Nbr (11–13). We set out to identify the functional *Ae. aegypti* Nbr ortholog by predicting all DEDDy-type 3’ - 5’ exonuclease domain-containing proteins, which were used in a phylogenetic analysis along with *Drosophila* Nbr (*DmNbr)*. This analysis identified AAEL005527 as a one-to-one ortholog of *DmNbr* (Figure 5A). In addition, a recent study verified that AAEL005527 exhibits Mn^2+^-dependent, ssRNA-specific 3’ – 5’ exonuclease activity (52). To evaluate the role of trimming for the formation of responder vpiRNA 3’ ends in *Ae. aegypti*, we combined AAEL005527/Nbr knockdown with SINV infection using the *gypsy*-targeted reporter viruses. Knockdown was efficient (92-93%, Figure S6A) and did not have a reproducible effect on viral RNA levels (Figure S6B). Aside from its role in piRNA 3’ end formation, Nbr is required for trimming of microRNAs (miRNAs), including miR-34-5p (53, 54). Thus, to verify that AAEL005527 is indeed the functional orthologue of *Drosophila* Nbr, we first assessed the effect of its depletion on trimming of two miRNA with heterogeneous 3’ ends in *Ae. aegypti*: miR-34-5p and miR-184 (55, 56). Indeed, knockdown of AAEL005527 resulted in a marked reduction of miR-34-5p trimming. Similarly, AAEL005527 knockdown resulted in a specific decrease of smaller miRNA-184 isoforms (Figure 5B – Northern blot 1), confirming that AAEL005527 is indeed the functional ortholog of *Drosophila* Nbr. Next, we analyzed the effects of Nbr and Zuc knockdown on viral responder piRNA size. Similar to our previous findings, knockdown of Zuc resulted in an electrophoretic shift of viral responder piRNAs towards higher sizes (Figure 5B – Northern blot 2). Interestingly, for all viruses tested, Nbr knockdown resulted in a reduction, specifically of shorter (<27 nt), responder piRNA isoforms, without affecting the larger isoforms (Figure 5B – Northern blot 2, Figure S6C). A similar reduction of shorter piRNA isoforms upon Nbr knockdown has previously been observed in *Drosophila* (12, 13). These findings suggest that mosquito Nbr trims pre-piRNAs generated through a Zuc-mediated endonucleolytic cut, but that only a minor fraction of such pre-piRNAs undergo trimming.

**Figure 5.**
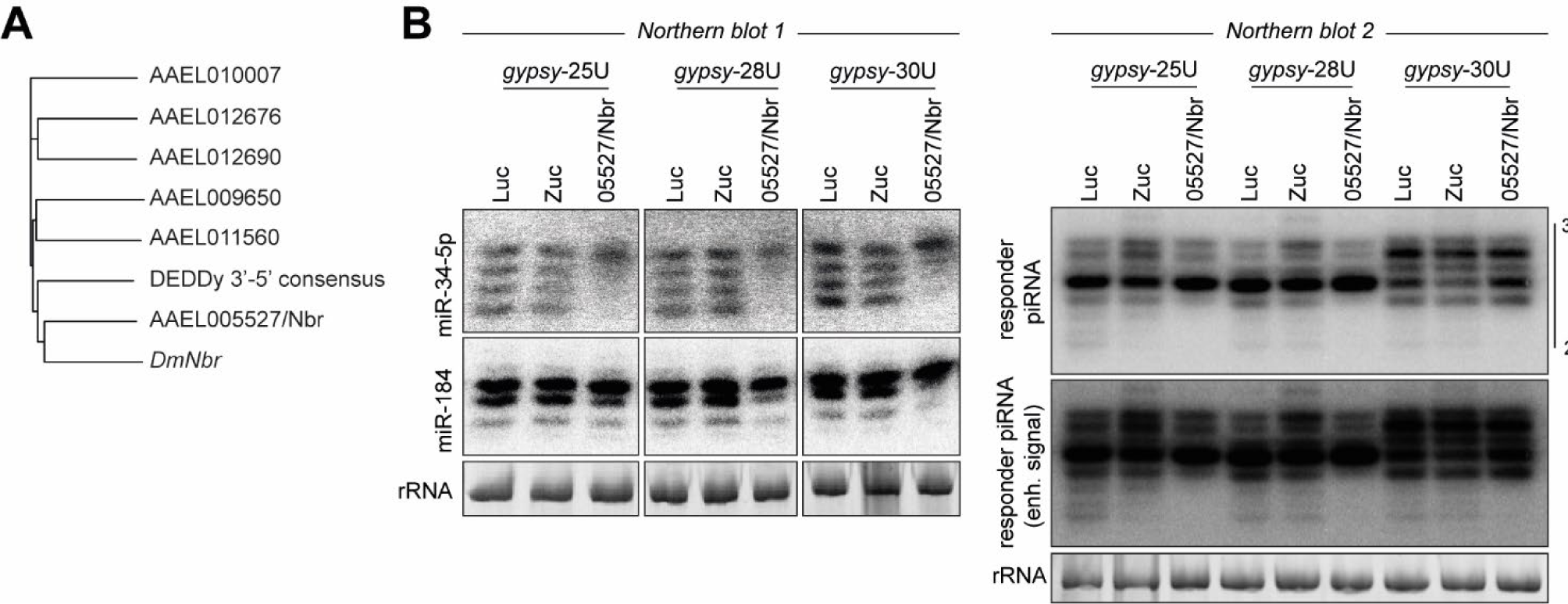
A subset of responder piRNAs undergoes Nbr-mediated trimming. **(A)** Phylogenetic tree based on the DEDDy 3’ – 5’ exonuclease domains identified in *Ae. aegypti*, along with the DEDDy consensus sequence and the DEDDy domain of *DmNbr*. **(B)** Northern blot analyses of miR-34-5p and miR-184 (left) and the viral responder piRNA (right) in Aag2 cells upon knockdown of Zuc and Nbr or control knockdown (Luc). RNA from the same knockdown experiment was analyzed on two separate northern blots. Ethidium bromide stained rRNA serves as a loading control. The Nbr knockdown phenotype has been observed in four independent infections.

The 3’ - 5’ exonucleases PNLDC1 and PARN-1 are responsible for trimming of piRNA 3’ ends in *B. mori* and *C. elegans*, respectively (57, 58). While PNLDC1 is not conserved in *Ae. aegypti* (11), a clear mosquito ortholog of PARN-1 can be identified: AAEL001426. Knockdown of this gene however, had no effect on responder piRNA 3’ end formation in our viral reporter system (Figure 4D, Figure S5C). In summary, we identified *Aedes aegypti* Nibbler to be involved in trimming of piRNAs and miRNAs.

### Targeting by an endogenous piRNA triggers trailer piRNA production

While endogenous piRNAs in *Ae. aegypti* show strong signatures of phased piRNA production (6), it is unknown whether RNA from cytoplasmic viruses is processed similarly through piRNA phasing. This is especially interesting as the genomes of *Ae. aegypti* and *Ae. albopictus* mosquitoes contain a high number of endogenous viral elements (EVEs). These non-retroviral sequence elements are enriched in piRNA clusters and, accordingly, give rise to abundant piRNAs (28,31–33), which may guide the slicing of cognate RNA from acutely infecting viruses. It has recently been shown that EVE-derived piRNAs are indeed able to target and inhibit newly infecting viruses (34, 35), yet, it remains unknown whether piRNA phasing can expand the vpiRNA sequence repertoire after an initial cleavage by an endogenous piRNA.

We first confirmed that we could detect signatures of piRNA phasing in our small RNA deep-sequencing data. In line with prior findings (6), we observed that the distance between transposon-derived piRNA 5’ ends was regularly phased in intervals of approximately 30 nt (Figure 6A). Strikingly, a similar periodicity was observed when we analyzed SINV-derived piRNAs (Figure 6B), indicating that viral RNA is also subjected to piRNA phasing. We noted that, compared to transposon-derived piRNAs, the periodicity of phasing of vpiRNAs is noisier after the second interval. This is likely explained by technical limitations of analyzing phasing signatures on the relatively small sequence of the SINV genome (approximately 11 kB in size).

**Figure 6.**
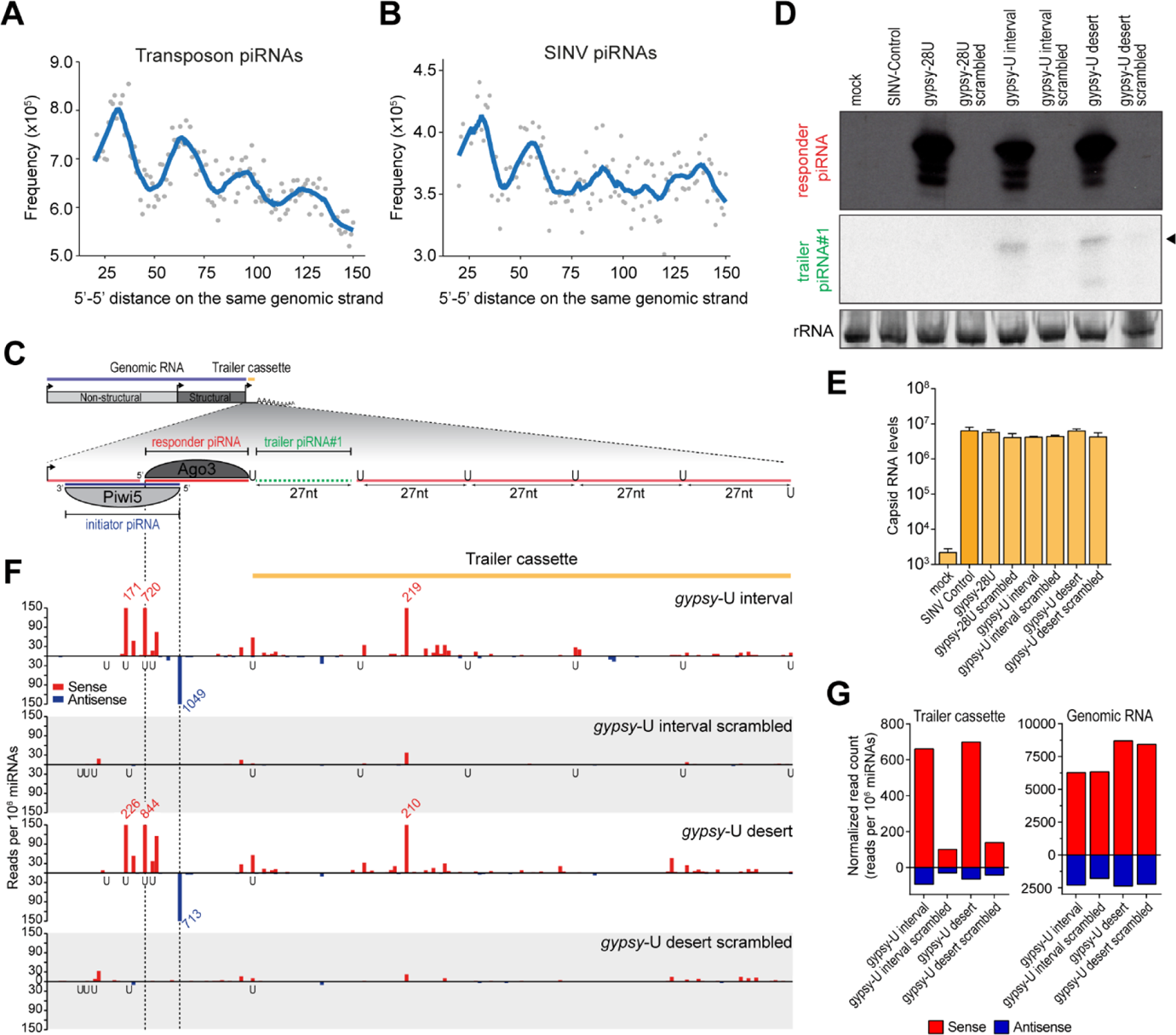
Targeting of viral RNA by an endogenous piRNA triggers trailer piRNA production. **(A-B)** Frequency distribution of piRNA 5’–5’ end distances for transposon (A) and SINV-derived (B) piRNAs. For each piRNA 5’ end, the number of piRNAs 5’ ends is counted in a downstream window of 20 to 150 nt. The combined distance frequencies are plotted and the curve is smoothened with local regression (LOESS). To reduce noise, only 5’ ends with a minimal read count of twenty were included in the analysis. **(C)** Schematic overview of the *gypsy*-U interval reporter virus. The inset shows a magnification of the non-coding reporter RNA expressed under control of the duplicated subgenomic promoter. This reporter RNA contains a target site for a Piwi5-bound *gypsy*-derived initiator piRNA, which guides the production of an Ago3-associated responder piRNA. The downstream sequence makes up the trailer cassette and either contains regularly spaced uridine residues (*gypsy-*U interval) or is devoid of uridine residues (*gypsy-*U desert). The responder piRNA and the first trailer piRNA are indicated in red and green, respectively. **(D)** Northern blot analysis of the responder and the first trailer piRNA (indicated by an arrowhead) produced from the indicated viruses. As a control, the target site was scrambled to abolish targeting by the *gypsy*-derived piRNA. The remainder of the responder piRNA site and the trailer cassette are identical to the respective non-scrambled U interval and U desert viruses. As additional controls, a virus bearing an intact target site, but no trailer cassette (gypsy-28U), and a virus that contains no insert (SINV Control) were used. The general structure of the *gypsy*-U interval virus is shown schematically in (C). rRNA stained with EtBr serves as a loading control. **(E)** RT-qPCR analyses of viral capsid RNA levels in Aag2 cells infected with the indicated viruses. Bars and whiskers show the mean and SD of three independent biological replicates. Unpaired two tailed t-tests with Holm-Sidak correction were used to determine statistically significant differences compared to SINV Control. **(F)** Normalized counts (in reads per 10^6^ miRNAs) of piRNA 5’ ends mapping to the initiator piRNA target site and trailer cassette of the indicated viruses. 5’ ends of initiator and responder piRNAs are indicated by dashed lines. Numbers in red and blue indicate counts that exceed the range of the y-axis. The position of uridine residues on the sense strand is indicated below the x-axis. Per condition, a single small RNA sequencing library was analyzed (same for G-H)). **(G)** Total number of normalized sense (red) and antisense (blue) piRNA-sized (24-33 nt) reads mapping to the trailer cassette (left) and genomic RNA (right) of the indicated viruses. The areas of virus denoted as the trailer cassette and genomic RNA are shown in yellow and blue in (C).

Since vpiRNAs were biased for downstream uridines (Figure 1D, S1B) and positioning of uridine residues directed the 5’ end formation of the first trailer piRNA in our reporter viruses (Figure 2C), we next assessed the contribution of uridine residues to piRNA phasing. To this end, we introduced an additional non-coding RNA sequence downstream of the *gypsy* and EVE initiator piRNA target sites, which we termed the trailer cassette. To direct sequential Zuc-mediated endonucleolytic cleavage, this cassette contained uridine residues at regularly spaced 27 nt intervals in an RNA sequence that was otherwise devoid of uridines (U interval viruses, schematically shown in Figure 6C and Figure S7C). As a control, these uridines were replaced by adenosine residues to create a trailer cassette completely devoid of uridines (U desert viruses). In Aag2 cells infected with these reporter viruses, but not control viruses in which the initiator piRNA target site was scrambled, responder piRNAs are abundantly produced (Figure 6D and Figure S7A). In cells infected with the *gypsy*- and EVE-U interval virus, we also detected the first trailer piRNA using northern blotting (Figure 6D, S7A). Interestingly, we also observed the production of the first trailer piRNA in cells infected with the U desert viruses (Figure 6D, S7A), suggesting a downstream uridine residue was not essential for the maturation of the 3’ end of the first trailer piRNA. Importantly, in cells infected with viruses lacking the entire trailer cassette (*gypsy*-28U or EVE25/28U), responder piRNAs but no trailer piRNAs are detected, indicating that trailer piRNAs are specifically derived from the trailer cassette (Figure 6D, S7A). Moreover, no trailer piRNAs were produced in cells infected with control viruses containing a scrambled target site (Figure 6D, S7A), indicating that trailer piRNA production depends on initial targeting by the endogenous piRNA. Of note, differences in piRNA abundance were not the result of changes in viral RNA levels, which was similar for all viruses used (Figure 6E, S7B).

To assess phased piRNA biogenesis beyond the first trailer piRNA, we sequenced small RNAs produced in Aag2 cells infected with interval and desert viruses, as well as their respective scrambled control viruses. Mapping piRNAs onto the trailer cassette reveals production of additional piRNAs in cells infected with the *gypsy* and EVE initiator piRNA targeted viruses (Figure 6F, Figure S7D). In contrast, barely any piRNAs mapping to the trailer cassette were recovered in cells infected with viruses bearing a scrambled target site (Figure 6G, left panel, Figure S7E, left panel). Viral piRNA production from the SINV genome upstream of the artificial reporter and trailer cassettes was unaltered (Figure 6G, right panel, Figure S7E, right panel), indicating that there are no differences in sensitivity of these viruses for processing by the vpiRNA biogenesis machinery. As both the pattern and level of piRNA production is highly similar between U interval and U desert viruses (Figure 6F-G, left panel, Figure S7D-E, left panel), the presence of uridine residues to guide Zuc-mediated endonucleolytic cleavage appears to be dispensable for trailer piRNA production in the context of the artificial trailer cassette. Importantly, these data show that initial targeting by a genome-encoded piRNA results in the production of additional piRNAs downstream of the target site, resulting in diversification of the viral piRNA pool.

### Conclusion

Altogether, our results indicate that during acute infection with a cytoplasmic RNA virus, endogenous piRNAs can initiate piRNA production from viral genomic RNA via the ping-pong amplification loop (Figure S8). The endonucleolytic and exonucleolytic activities of Zuc and Nbr are involved in maturation of the 3’ ends of piRNAs. Importantly, cleavage of viral RNA by an endogenous piRNA triggers the diversification of piRNA sequence repertoire by triggering the production of additional trailer piRNAs from the downstream cleavage fragment. These findings indicate that a few cleavage events by individual genome-encoded piRNAs are sufficient to launch a piRNA response that may eventually become independent of an endogenous trigger.

## DATA AVAILABILITY

Small RNA sequencing data has been deposited to the NCBI Sequence Read Archive under accession number SRP272125.

## ACKNOWLEDGEMENTS

We thank the members of the laboratory for critical discussion of this manuscript and Bas Pennings for his help with cloning recombinant Sindbis virus constructs. We acknowledge the Carnegie Institute for providing the Drosophila Gateway Vector Collection.

## Supplemental text

### An EVE-derived piRNA has the potential to trigger responder piRNA production

Besides the *gypsy* initiator piRNA triggered reporter virus described in the main text, we generated a second set of viruses bearing target sites for an abundant Piwi5-associated piRNA derived from an EVE of flaviviral origin (described as FV53 in (1), Figure S2A, C). As the target site for the EVE-derived initiator piRNAs can be recognized by two piRNA isoforms that align at their 3’ end and differ 3 nt in size (27 and 30 nt), target cleavage may result in the production of two Ago3-bound responder piRNAs isoforms that have an offset of 3 nt (blue shaded inset in Figure S2A).

Similar to the *gypsy* target site bearing viruses (Figure 2B), responder piRNA 3’ end sharpness is moderately reduced as a function of increased distance between the initiator piRNA cleavage site and the first downstream uridine residue (Figure S2D), suggesting a role for exonucleolytic trimming in responder piRNA maturation.

For all viruses tested here, we observe abundant responder piRNAs as well as the production of a first trailer piRNA (indicated with yellow arrowheads in Figure S2E). As the trailer piRNA 5’ ends perfectly overlap with the uridine residue, we propose that this residue guides cleavage of the viral RNA, simultaneously generating the responder pre-piRNA 3’ end and the trailer piRNA 5’ end, as also observed for the *gypsy* targeted viruses (Figure 2E).

Aside from these responder and trailer piRNAs, additional abundant sense piRNAs are produced from the sequence upstream of the initiator piRNA slice site (Figure S2E), suggesting that other, partially complementary, initiator piRNAs may target the sequence and instruct production of these additional sense piRNAs.

**Figure S1.**
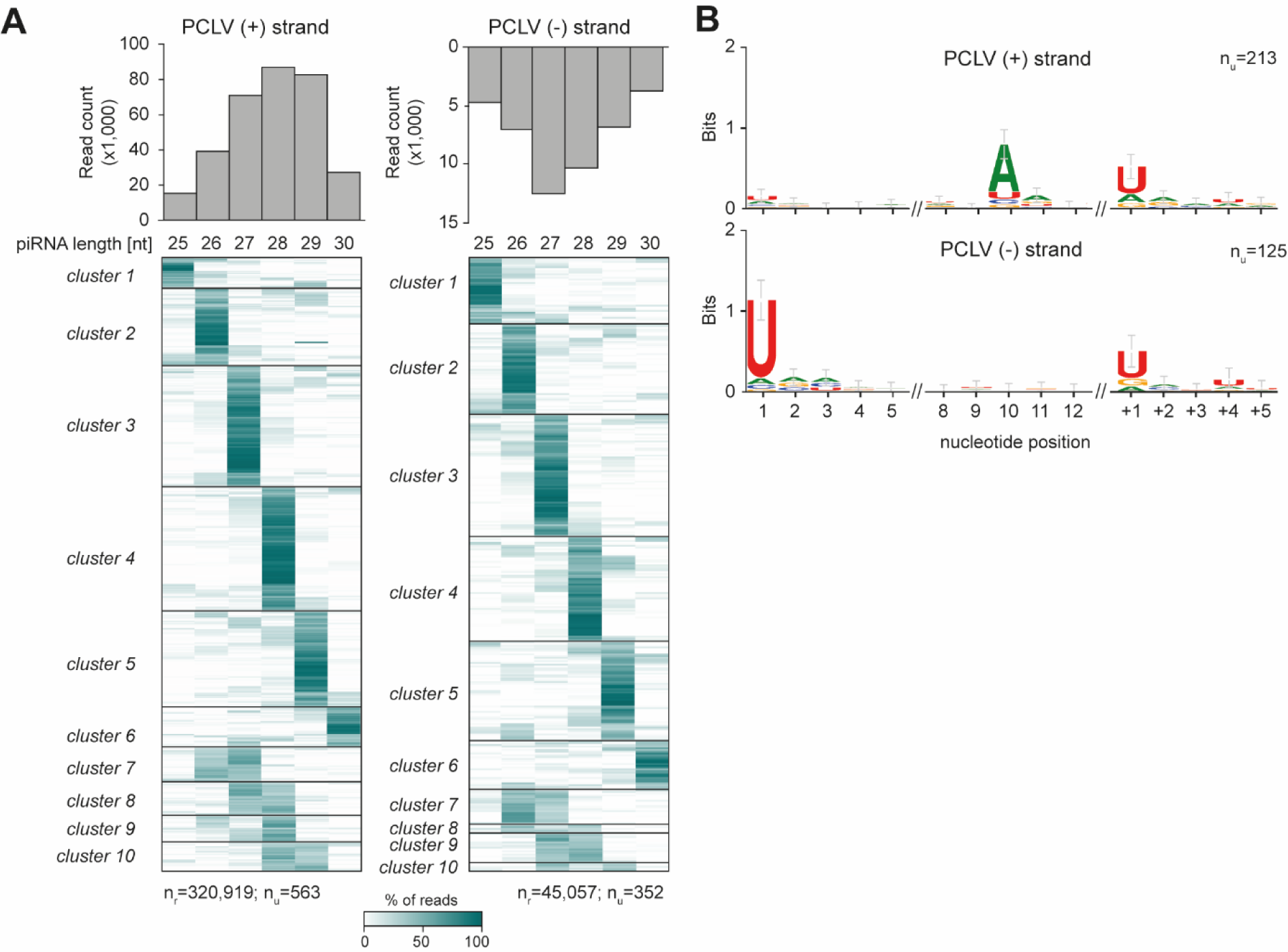
PCLV-derived piRNAs have sharp 3’ ends (A) Analysis of lengths of individual piRNAs derived from Phasi Charoen-like virus (PCLV). Data were generated from the combined three dsLuc small RNA libraries published in (2). Size profiles of piRNAs mapping to the sense ((+), left) and antisense ((-), right) strand of the virus genome are shown in the top panels. The heat maps show the relative size distribution of PCLV piRNAs that share the same 5’ end (one 5’ end is one line in the heat map). Shades of green indicate the percentage of reads contributing to the indicated piRNA length, white represents absence of reads of the specific size. The number of reads (nr) and the number of unique piRNAs 5’ ends (nu) that underlie the heat map are indicated. A minimum of 20 reads per unique piRNA position was required to be included in the analysis. **(B)** Nucleotide biases for the indicated nucleotide positions of PCLV-derived vpiRNAs and the genomic region downstream (+1 to +5) of the vpiRNAs. Only vpiRNAs from (A) that had a dominant length (supported by at least 75% of reads) were considered in this analysis and all reads were collapsed to unique sequences, irrespective of read count. nu indicates the number of sequences underlying the sequence logo.

**Figure S2.**
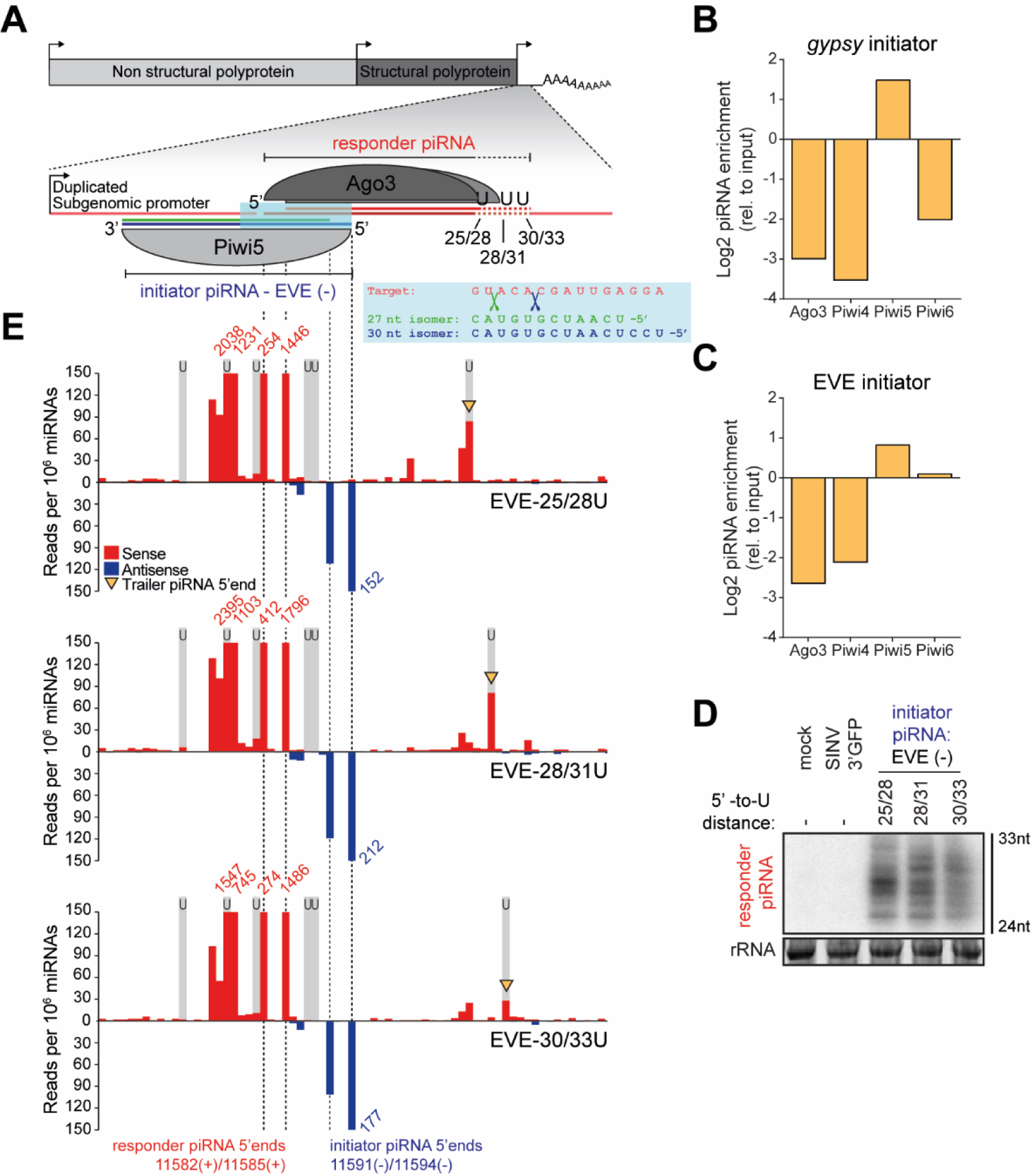
An EVE-derived piRNA triggers responder piRNA production from an acutely infecting RNA virus (A) Schematic depiction of the SINV-based viral reporter system in which responder piRNA production is triggered by EVE initiator piRNAs. As the Piwi5-associated EVE initiator piRNA is expressed as two isoforms that align at their 3’ end (blue and green lines), endonucleolytic cleavage may generate two Ago3-bound responder piRNAs (red lines) with a 5’ end offset of 3 nt . The blue shaded inset depicts an enlargement of a part of the viral target RNA (red), the 5’ ends of the two EVE-piRNA isoforms (27-nt: green and 30-nt: dark blue); the green and dark blue scissors represent their respective slice sites. **(B-C)** Log2 fold enrichment of the *gypsy* (B) and EVE (C) initiator piRNAs in PIWI protein immunoprecipitation (IP) small RNA libraries. Per condition, a single sequencing library previously described in (3, 4) was analyzed. **(D)** Northern blot analyses of responder piRNAs produced in Aag2 cells infected with indicated EVE reporter viruses. The distance between responder piRNA 5’ ends and the first downstream uridine residue is indicated. 24 and 33 nt size markers are inferred from EtBr staining of a small RNA marker, and EtBr stained rRNA serves as loading control. The northern blot is representative of three independent biological replicates. **(E)** Visualization of 5’ ends of sense (red) and antisense (blue) piRNAs (24-33 nt) mapping to the non-coding reporter RNA sequence. Predicted initiator and responder piRNA 5’ ends are indicated by dashed lines and positions of uridine residues on the sense strand are denoted by light grey shading. Red and blue numbers indicate read counts at positions where they exceed the y-axis range and yellow arrowheads indicate 5’ ends of putative trailer piRNAs. A single sequencing library was analyzed for each virus infection.

**Figure S3.**
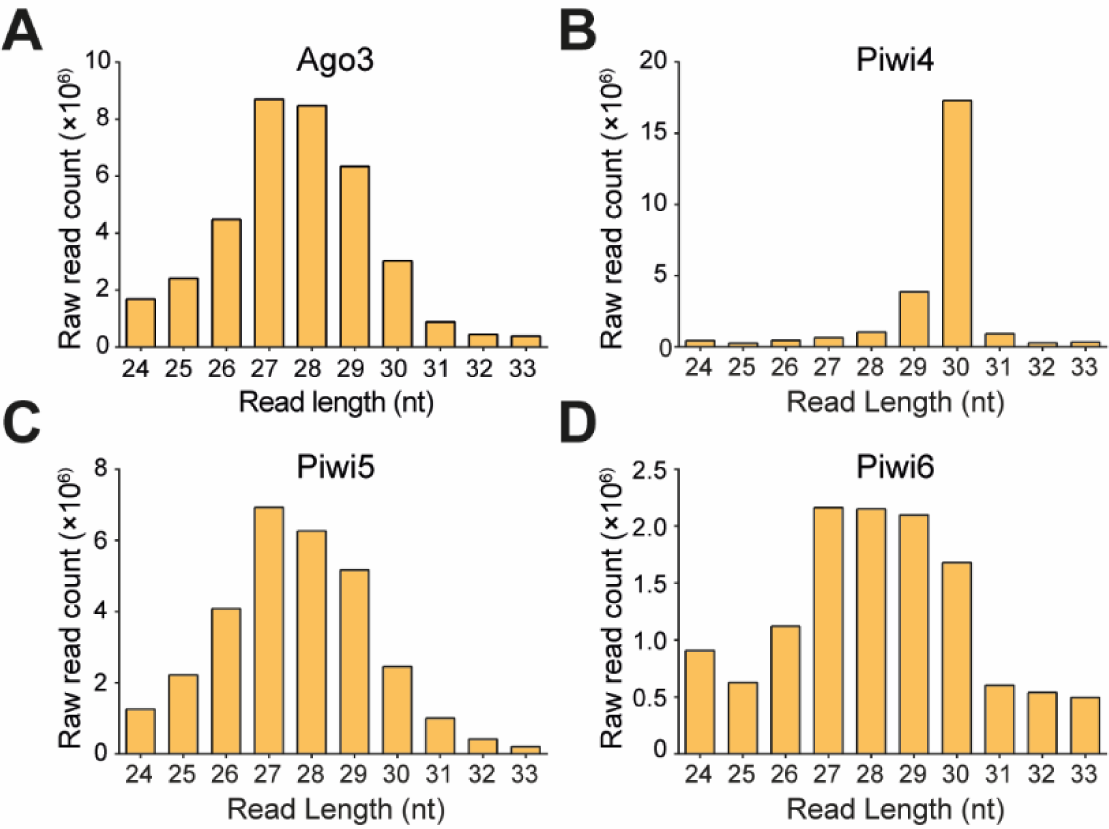
Size profiles of piRNAs associated with somatic PIWI proteins (A-D) Size distribution of piRNA-sized (24-33 nt) small RNA reads in Ago3 (A), Piwi4 (B), Piwi5 (C), and Piwi6 IP (D) small RNA sequencing libraries. Piwi4 IP is dominated by two highly abundant piRNAs (tapiR1 and tapiR2 of 30 and 29 nt in size, respectively) that are involved in the degradation of maternally provided transcripts during embryonic development (4). Counts are raw, unmapped reads from our previously published small RNA sequencing data from endogenous PIWI protein IPs. (3, 4). For each IP, a single sequencing library was analyzed.

**Figure S4.**
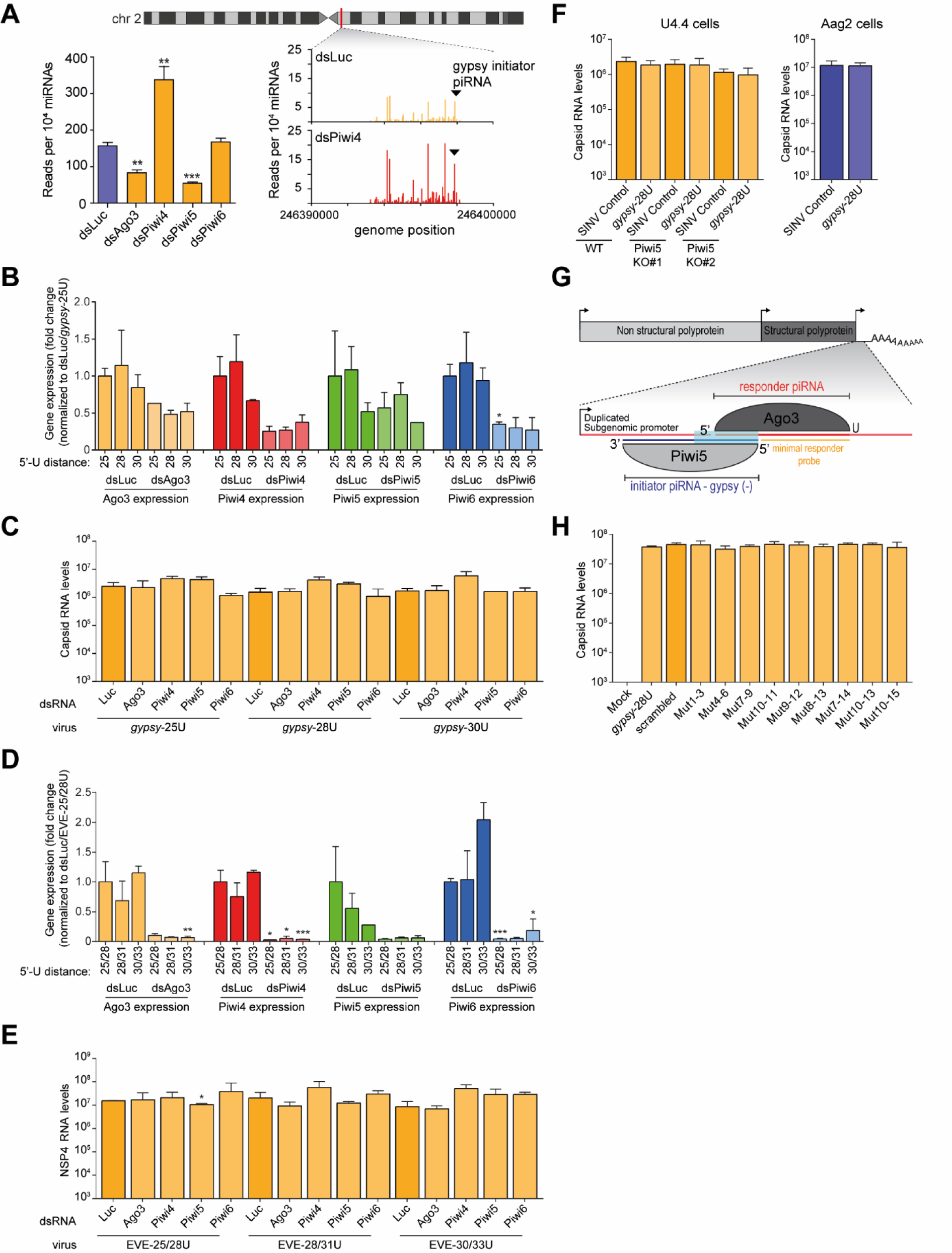
PIWI-knockdown does not affect virus replication (A) piRNA abundance at the genomic locus from which the gypsy initiator piRNA is derived. The bar chart indicates normalized piRNA counts in PIWI knockdown libraries compared to a control (dsLuc) knockdown published in (2). The line graph shows the genome distribution of piRNAs at chromosome 2 from position 246,390,000 to 246,400,000 for the control and Piwi4 knockdown datasets analyzed in the bar chart. The mean read count of thee libraries is shown. The black arrowhead indicated the position of the gypsy initiator piRNA that triggers responder piRNA production in the reporter viruses. **(B-C)** RT-qPCR analyses of relative PIWI gene expression (B) and subgenomic capsid RNA levels (C) in the samples used in Figure 3C. PIWI gene RNA levels are shown as a fold change relative to dsLuc treated cells infected with *gypsy*-25U. **(D-E)** RT-qPCR analysis of PIWI knockdown efficiency (D) and genomic NSP4 RNA levels in the samples used in Figure 3D. **(F)** RT-qPCR analyses of capsid RNA levels in the indicated U4.4 (yellow) and Aag2 (blue) samples used in Figure 3E. **(G)** Schematic representation of the *gypsy*-28U virus that was used to study the effect of target site mutations on responder piRNA production (Figure 3G-H). The sequence in which seed and slice site mutations were introduced is indicated by light blue shading and the ‘minimal responder’ probe used to detect responder piRNAs in the northern blot of panel 3H is indicated in yellow. This minimal responder probe hybridizes to the last 18 nt of the 3’ end of responder piRNAs, which are identical for all viruses. **(H)** RT-qPCR analysis of capsid RNA levels in samples used in Figure 3H. For all bar charts in this Figure, bars and whiskers show the mean and standard deviation (SD) of three independent biological replicates and unpaired two tailed t-tests with Holm-Sidak correction were performed for statistical analyses (* P < 0.05, ** P < 0.005, *** P < 0.0005).

**Figure S5.**
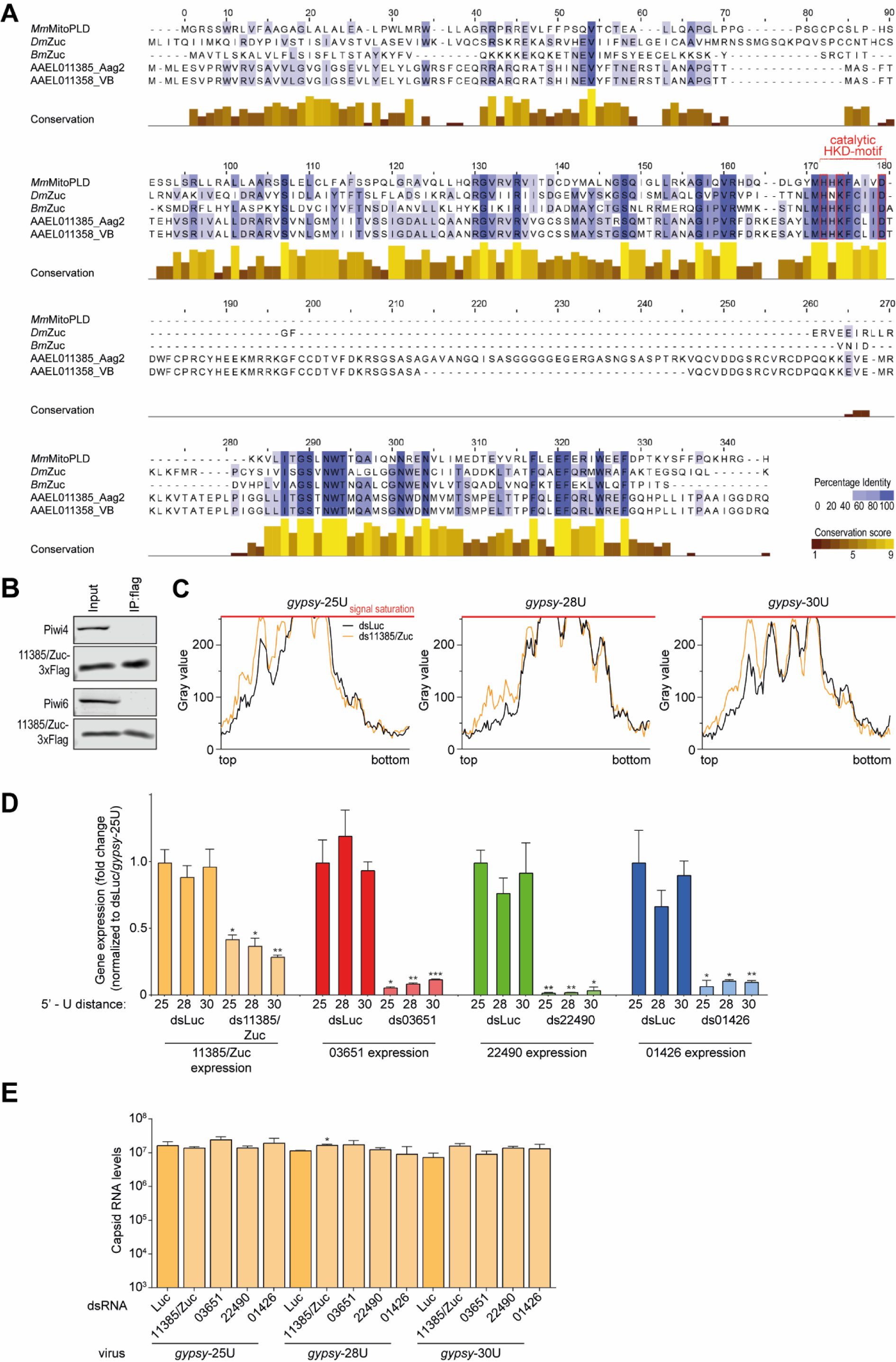
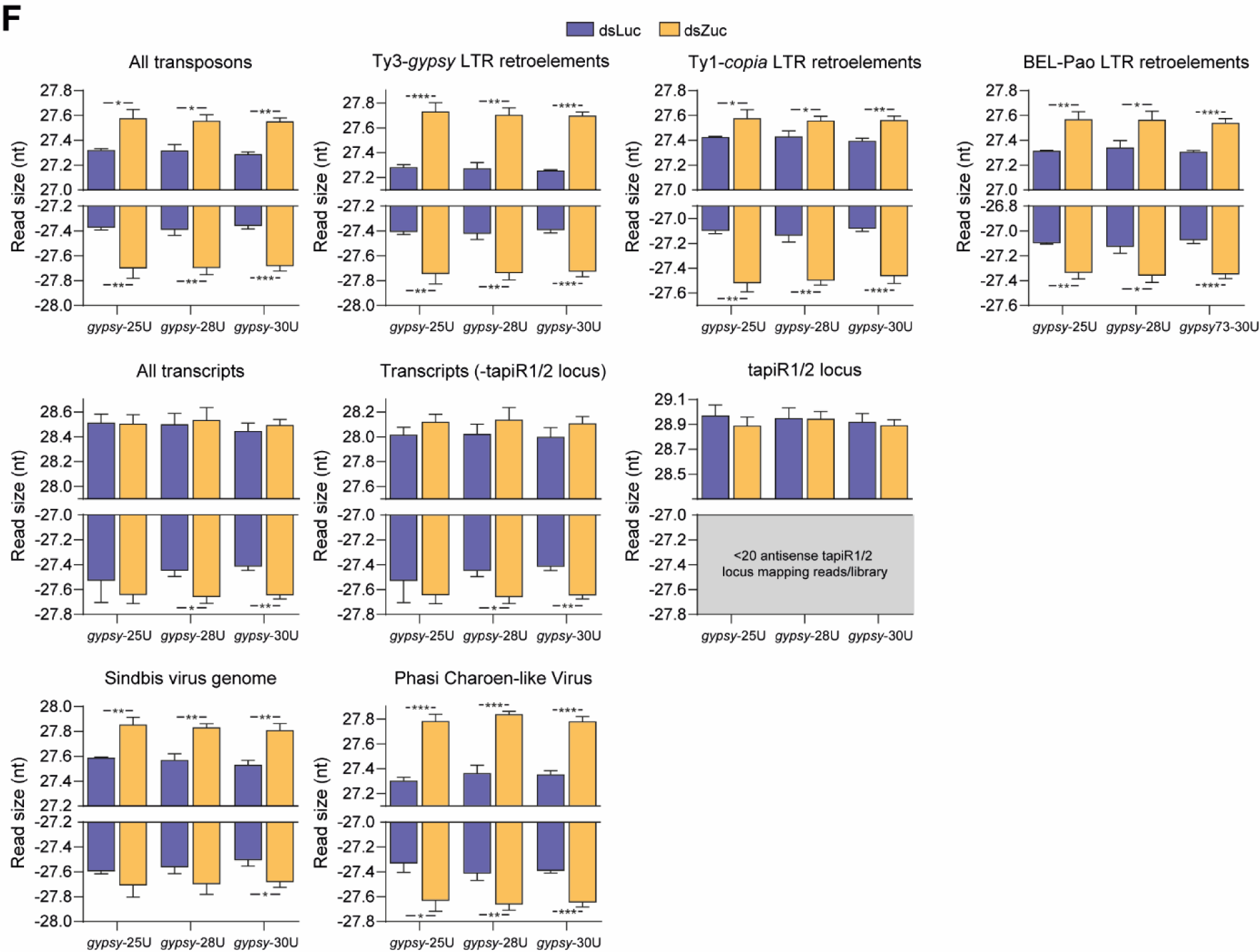
Mosquito Zuc is involved in 3’ end formation of piRNAs from various substrates (A) Multiple sequence alignment of Zucchini orthologs from *D. melanogaster* (*Dm*Zuc)*, B. mori* (*Bm*Zuc) and *M. musculus* (*Mm*MitoPLD), and AAEL011385 as annotated in VectorBase (VB) and as sequenced from Aag2 cells. Residues shaded in blue are shared between ≥3 of the proteins, and the brown-yellow bars indicate conservation of physicochemical properties at each position. **(B)** Western blot analyses of Piwi4 and Piwi6 in the same Zuc-3xflag-IP material that was used in Figure 4C. **(C)** Quantification of the responder piRNA signal from the northern blot in Figure 4D, analyzed for the three *gypsy*-derived piRNA targeted viruses separately. The X-axis represents the position on the northern blot from top to bottom, the Y-axis shows responder piRNA signal intensity. **(D-E)** RT-qPCR analyses of the knockdown efficiency of the indicated genes (D) and viral capsid RNA levels (E). For (D), gene expression levels are depicted as fold changes relative to dsLuc treated cells infected with the *gypsy*-25U virus. Knockdown efficiencies and viral RNA levels were tested against dsLuc treated samples infected with the same virus. **(F)** Average size of piRNAs (24-33 nt) derived from the indicated substrates in small RNA sequencing libraries of dsLuc and dsZuc treated Aag2 cells. The average length of two piRNAs (tapiR1/2) involved in degradation of maternal transcripts during embryogenesis (4), is not affected by Zuc knockdown. As virtually no antisense reads map to the tapiR1/2 locus, these data are not shown. For all bar charts, bars and whiskers represent mean and SD of three independent replicates. Unpaired two tailed t-tests with Holm-Sidak correction for multiple comparisons were used to determine statistical significance (* *P* < 0.05, ** *P* < 0.005, *** *P* < 0.0005).

**Figure S6.**
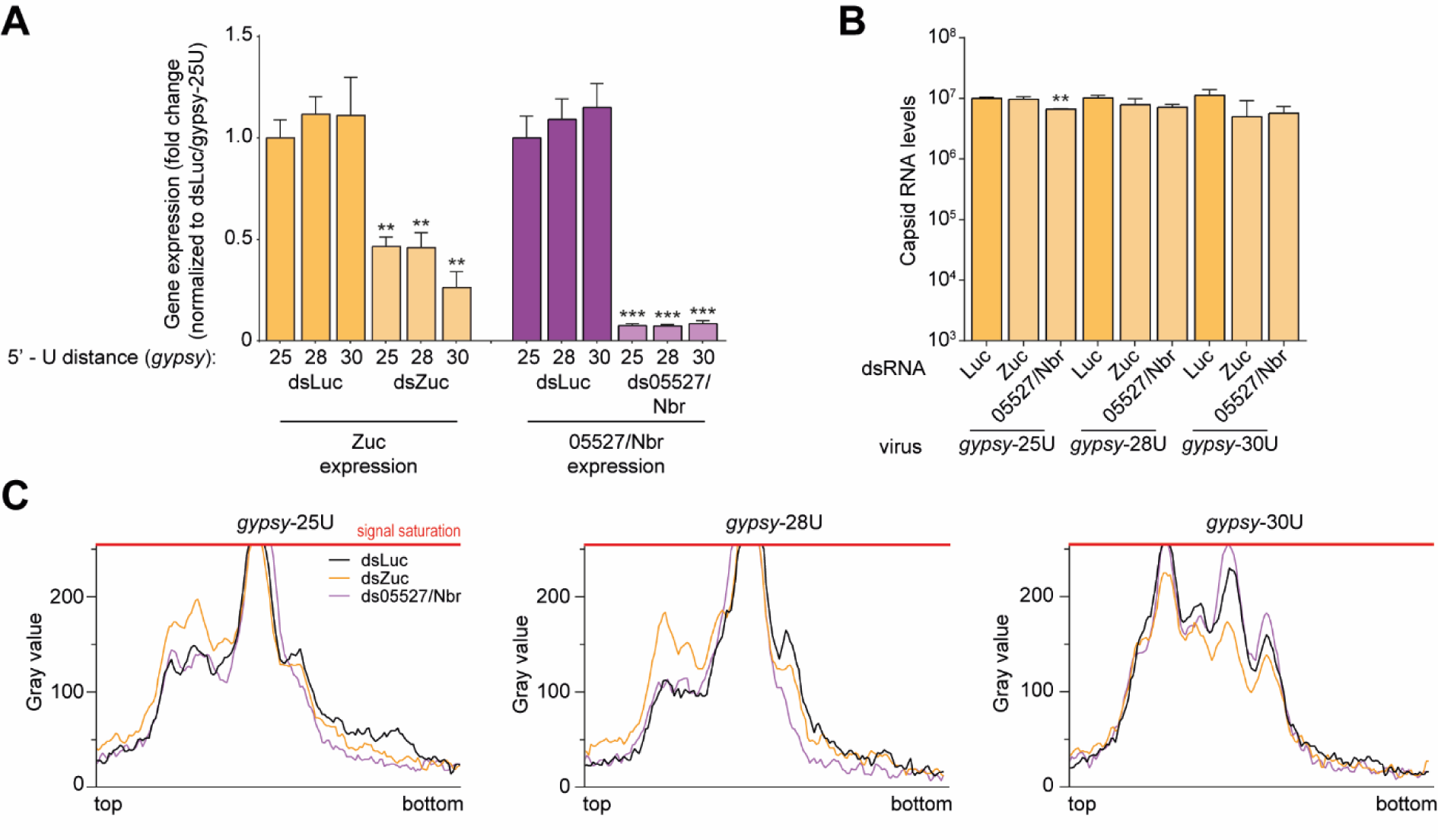
Nbr- and Zuc knockdown affect piRNA biogenesis (A-B) RT-qPCR analyses of the knockdown efficiencies of indicated genes (A) and viral capsid RNA levels (B). Bars and whiskers represent the mean +/- SD of three independent biological replicates. For (A), expression values represent the fold change relative to dsLuc treated cells infected with the *gypsy*-25U virus. Unpaired two tailed t-tests with Holm-Sidak correction for multiple comparisons were used to determine statistical significance (** *P* < 0.005, *** *P* < 0.0005). Knockdown efficiencies and viral RNA copy numbers were tested against dsLuc treated Aag2 cells infected with the same virus. **(C)** Quantification of the viral responder piRNA signal on northern blot 2 in Figure 5B, showing the effect of Nbr knockdown on responder piRNA size during infection with the indicated viruses. The X-axis represents the position on the northern blot from top to bottom, the Y-axis represents responder piRNA signal intensity.

**Figure S7.**
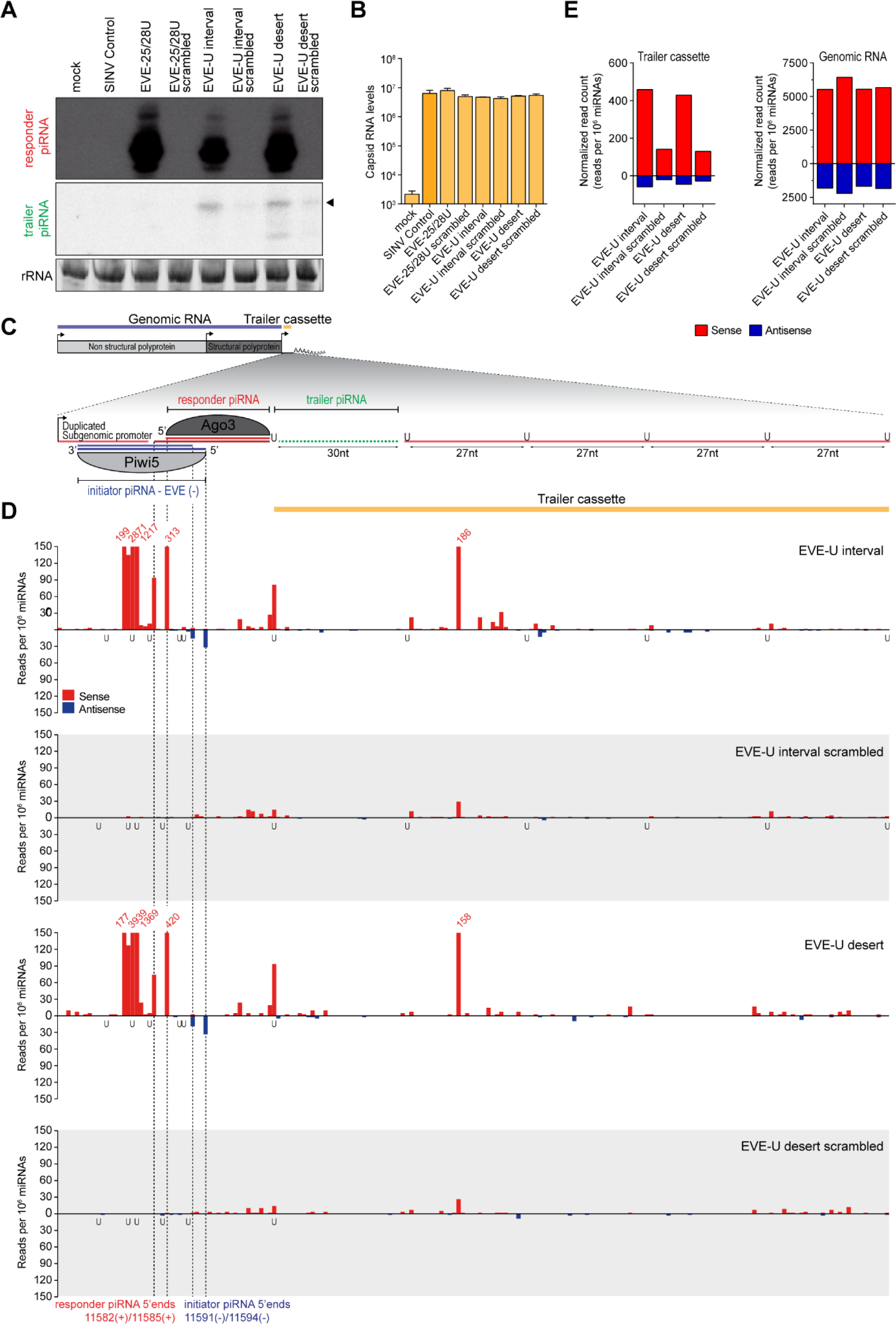
An EVE-derived piRNA triggers trailer piRNA production downstream of its target site (A) Northern blot analyses showing the production of the responder piRNA and the first putative trailer piRNA (indicated by an arrowhead) from the indicated viruses. As a control, the target site was scrambled to abolish targeting by the EVE-derived initiator piRNA. The remainder of the responder piRNA site and the trailer cassette was identical to the U interval and U desert viruses. Additional controls include a virus that lacks the trailer cassette, but contains an EVE piRNA-target site (FV25-28) and a control virus without insert (SINV Control). A schematic overview of the EVE targeted interval virus is shown in (C). rRNA stained by EtBr serves as loading control. n = one biological replicate. **(B)** Capsid RNA levels in Aag2 cells infected with indicated viruses as determined by RT-qPCR. Bars and whiskers denote the mean and SD of three independent biological replicates, respectively. Unpaired two tailed t-tests with Holm-Sidak correction were used to compare differences with the control virus. **(C)** Schematic overview of the EVE interval virus. In this virus, a duplicated subgenomic promoter drives the expression of a non-coding reporter RNA (shown in the magnification), which contains a target site for two isoforms of a Piwi5-associated EVE-derived initiator piRNA (shown in blue). Slicing of the reporter RNA in the ping-pong amplification loop thus may give rise to two differently sized Ago3-associated responder piRNAs (shown in red). Located downstream of this responder piRNA is the trailer cassette, which either contains uridine residues at regularly spaced intervals (EVE-U interval) or is completely devoid of uridines (EVE-U desert). The first trailer piRNA, which was detected in (A), is shown in green. **(D)** Normalized counts of 5’ ends of sense (red) and antisense (blue) piRNA-sized reads (24-33 nt) mapping to the non-coding reporter RNA. The 5’ ends of the two EVE-derived initiator piRNA isoforms, as well as the 5’ ends of corresponding responder piRNAs are indicated by dashed lines. The position of uridine residues is indicated below the x-axis. The numbers in red indicate normalized read counts that exceed the range of the y-axis. Per condition, a single small RNA sequencing library was analyzed (same for (E)).**(E)** Quantification of the number of sense (red) and antisense (blue) piRNAs mapping to the trailer cassette (left) and genomic RNA (right) of the indicated recombinant Sindbis viruses. The areas of the viruses denoted as trailer cassette and genomic RNA are indicated in (C) in yellow and blue, respectively.

**Figure S8.**
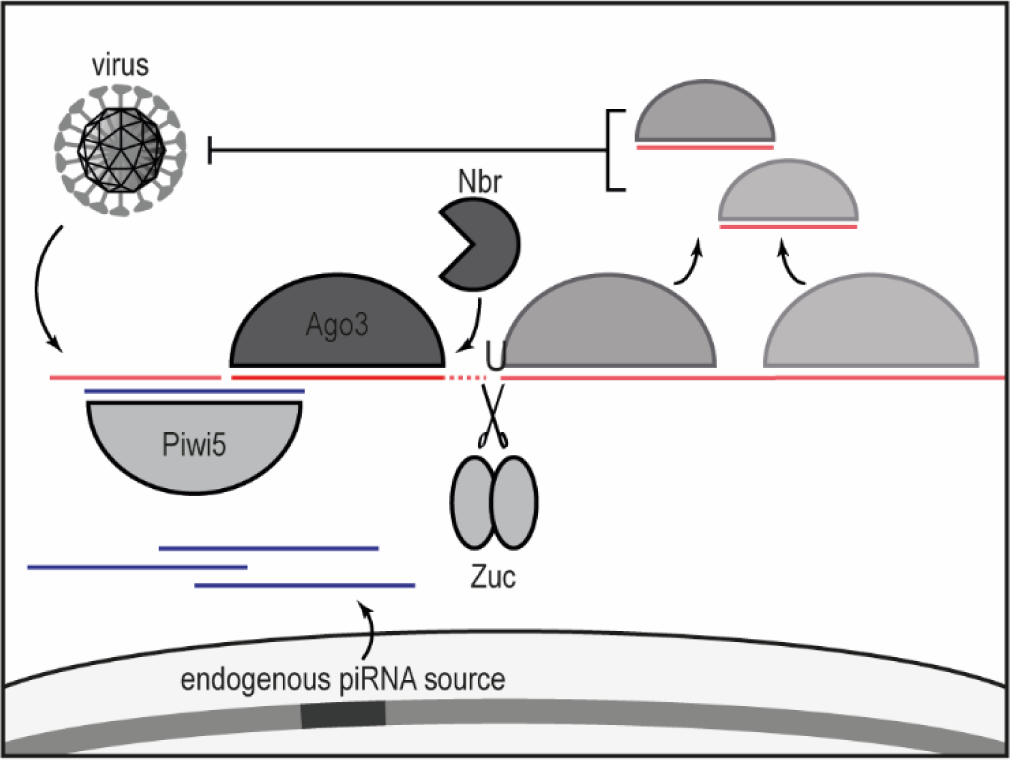
A model of piRNA-based adaptive immunity in mosquitoes. Genome-encoded endogenous viral elements (EVE) or other endogenous loci give rise to a pool of Piwi5-associated initiator piRNAs that have the potential to target newly infecting viruses. Upon infection with a virus containing a cognate sequence, EVE-derived piRNAs trigger the production of Ago3-bound responder piRNAs from the viral RNA, which are generated and maturated by the combined activities of Zuc and Nbr. In addition, targeting of the viral RNA by the ping-pong machinery initiates the processing of the cleavage fragment into additional downstream trailer piRNAs, thus expanding the piRNA sequence pool that is able to target viral RNA.

## Supplemental materials and methods

### Cell culture, dsRNA transfection and infection of Aag2 and U4.4 cells

*Ae. aegypti* Aag2 and *Ae. albopictus* U4.4 cells were grown in Leibovitz’s L-15 medium (Invitrogen) supplemented with 10% fetal bovine serum (Gibco), 50 U/ml Penicillin, 50 μg/mL Streptomycin (Invitrogen), 1x Non-essential Amino Acids (Invitrogen) and 2% Tryptose phosphate broth solution (Sigma) at 25°C. Cell lines were maintained by splitting twice weekly according to confluency.

For knockdown experiments, ∼1×10^6^ cells were seeded in 6-well plates and allowed to attach for 16-24 hrs. Subsequently, cells were transfected with gene-specific dsRNA using X-tremeGENE HP DNA Transfection Reagent (Roche) according to the manufacturer’s instruction, and re-transfected 48 hrs later to ensure sustained knockdown (750 ng dsRNA with 3 µL X-tremeGENE HP per well). Three hrs after the second transfection, cells were infected with indicated viruses at a multiplicity of infection (MOI) of 0.1 and RNA was harvested 72 hrs post infection. In experiments where no knockdown was performed, cells were infected 16-24 hrs after seeding at MOI = 0.1, followed by RNA extraction at 72 hrs post infection.

### Generation of reporter viruses

To introduce the target sites for *gypsy*- and EVE-initiator piRNAs, as well as the responder reporter cassette, the pTE3’2J-GFP plasmid encoding the previously described SINV-GFP (5, 6) was first digested using XbaI to remove the GFP-gene. Target sites and the reporter locus were subsequently introduced downstream of the duplicated subgenomic promoter by ligation of annealed oligonucleotides (see below), making use of the overhangs generated by the XbaI enzyme (initiator piRNA target site in red, downstream uridine position is indicated in **bold** font).

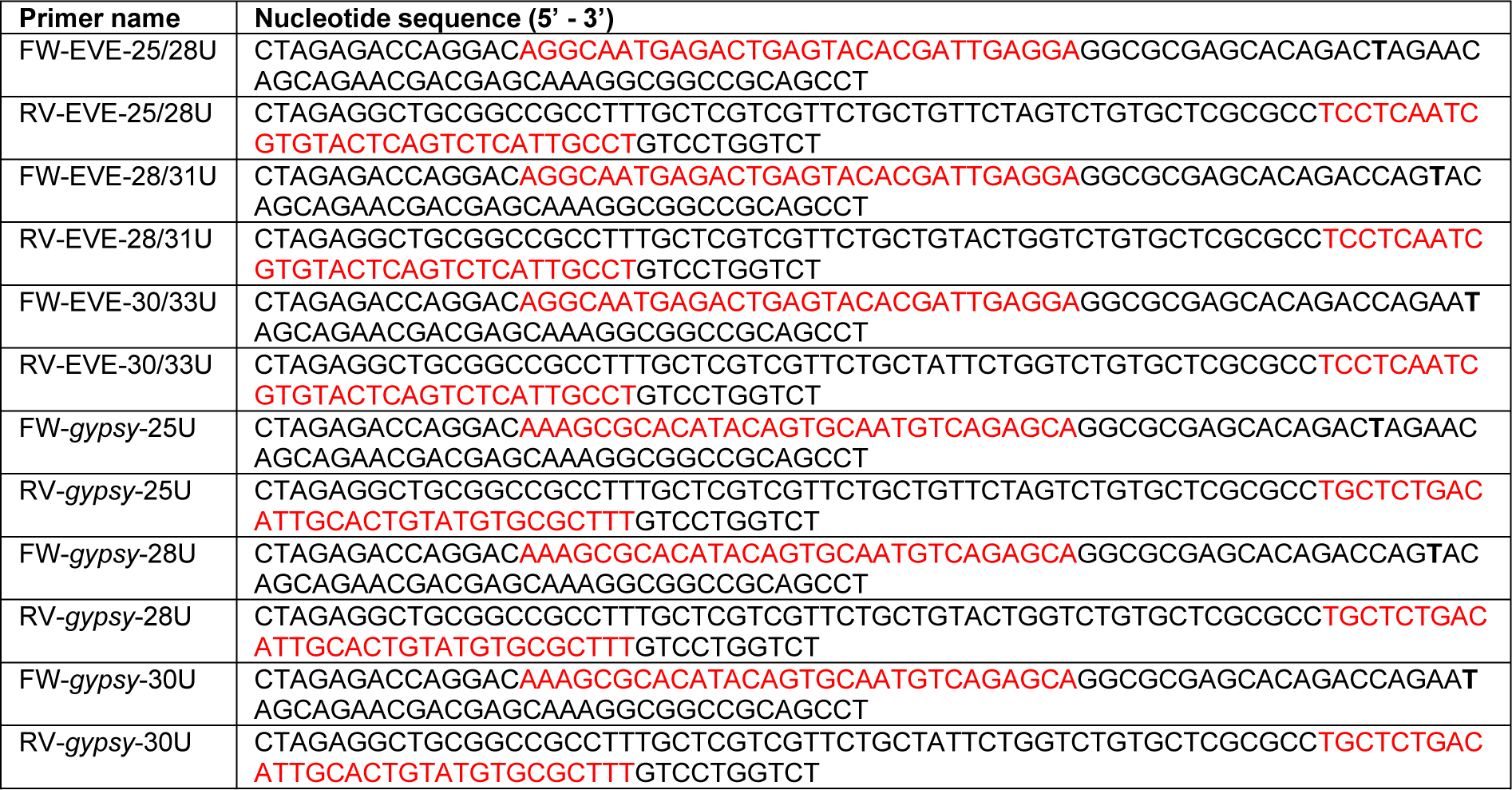

To introduce the U-interval and U-desert reporter cassettes, the *gypsy*-28U and EVE-25/28U viruses were digested using NotI followed by ligation of annealed oligo’s (see below).

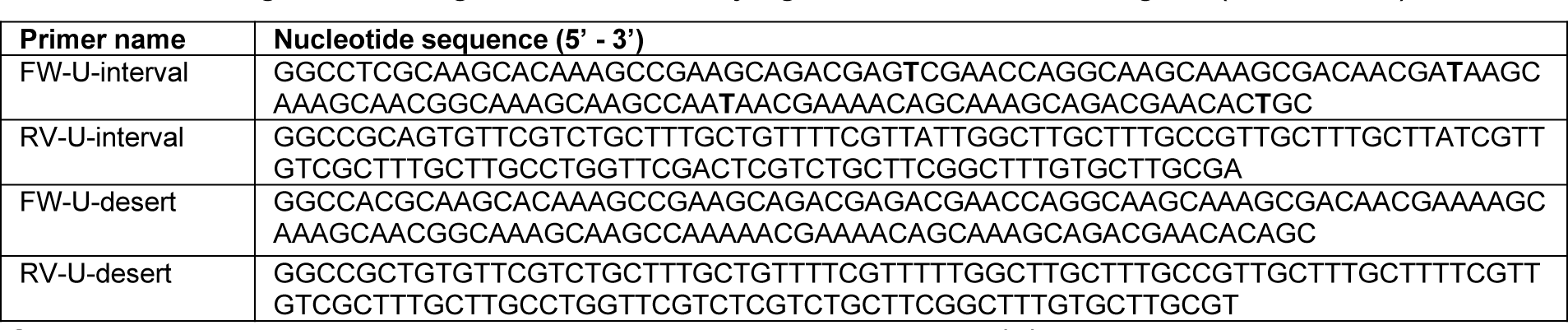

Subsequently, viruses were grown as described previously (7).

### Generation of target site mutant viruses

Target site mutations were introduced into the plasmid encoding the *gypsy*-28U virus by mutagenesis PCR, using the primers shown below. PCR products were DpnI-treated and In- fusion (Takara Biotech) was used to circularize the plasmid for transformation. After verification of the sequence by Sanger sequencing, viruses were grown as described previously (7).

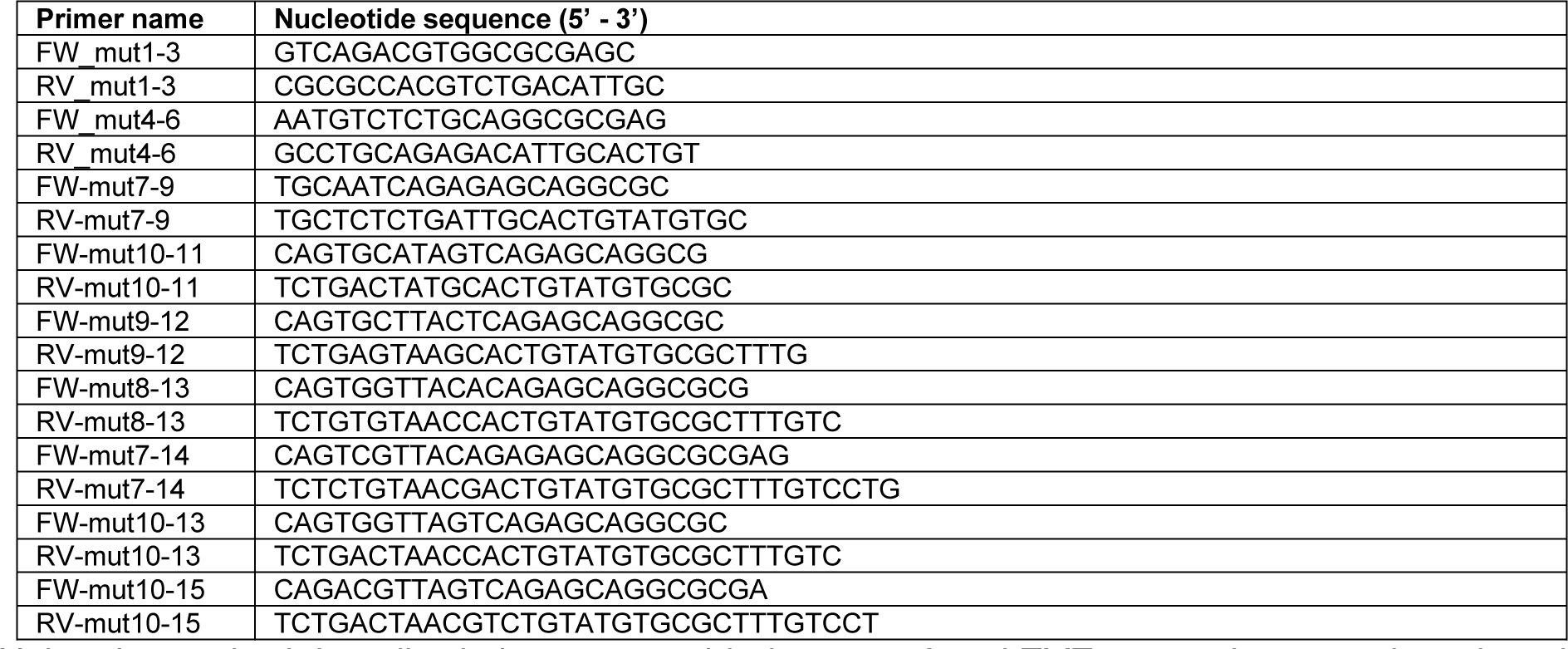

Using the method described above, scrambled *gypsy 73* and EVE target sites were introduced into the *gypsy*-28U, *gypsy*-U interval, *gypsy*-U desert, EVE-25/28U, EVE-U interval and EVE- U desert backbones, using primers listed below.

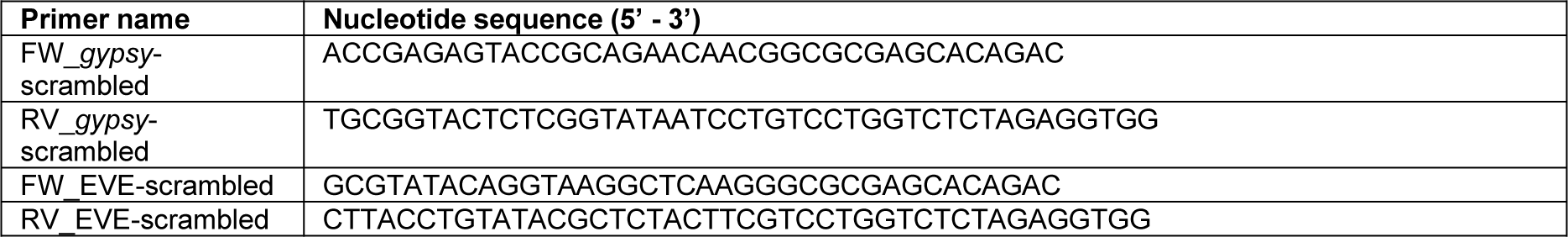

### dsRNA production for knockdown experiments

Gene-specific PCR products bearing T7 promoter sequences at both ends were generated in one of two ways. Either the T7 promoter sequence was introduced directly with the gene-specific PCR, or a universal GC-rich tag was introduced in the first PCR, to which the T7 promoter sequence was added in a second PCR. Oligonucleotides used to create the T7- promoter tagged PCR products are shown below (T7 promoter sequence in **bold** font, the universal GC-rich tag in underlined font):

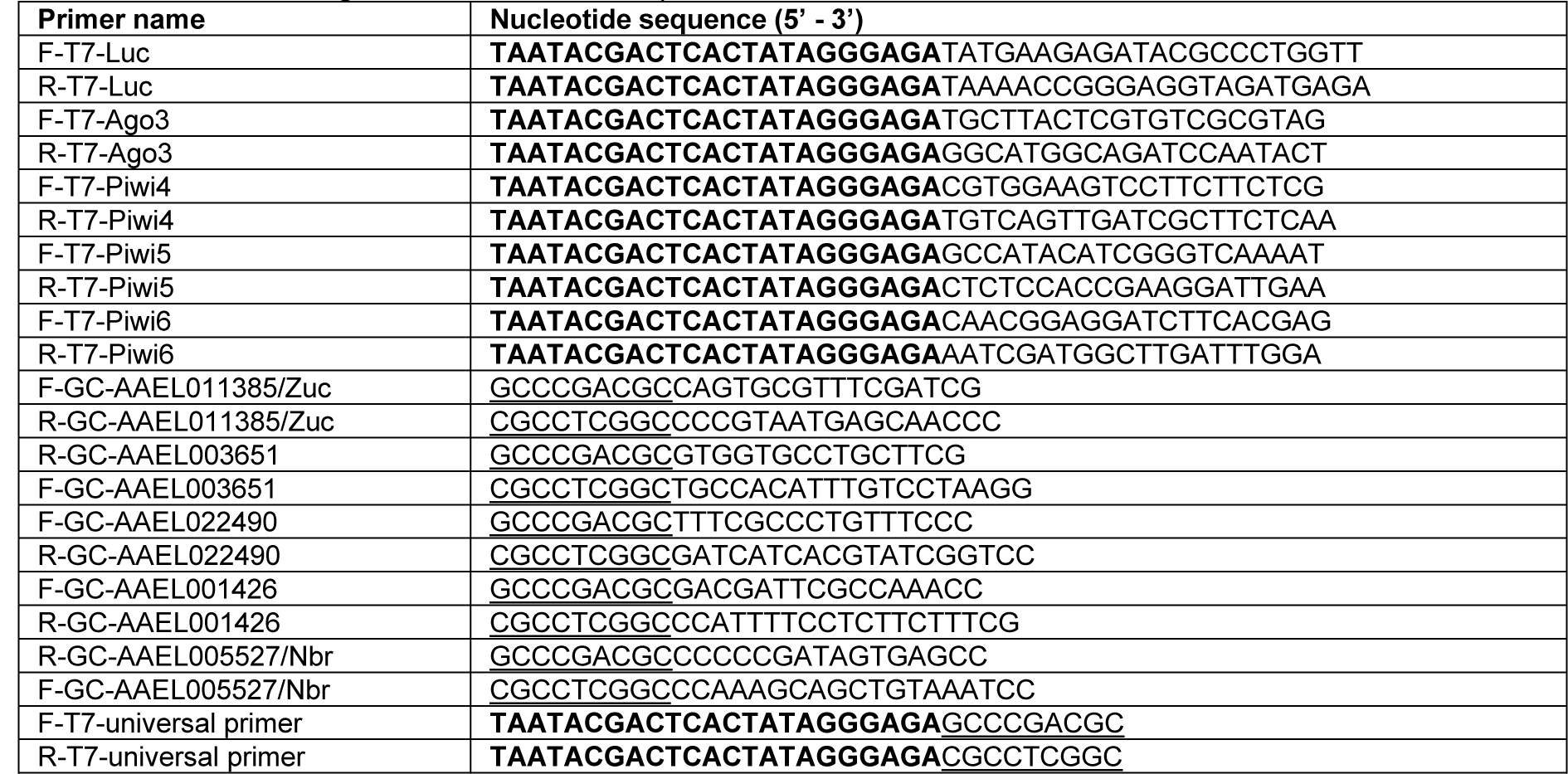

Subsequently, these PCR products were used as template for *in vitro* transcription by T7 polymerase for 4 hours at 37°C. Afterwards, the *in vitro* transcribed RNA was heated to 80°C and gradually cooled to room temperature to allow the formation of dsRNA, which was subsequently purified using the GenElute Mammalian Total RNA kit (Sigma).

### Generation of Piwi5 knockout U4.4 cells

Description and characterization of Piwi5 knockout U4.4 cells is described elsewhere (manuscript in prep).

### Phylogenetic analyses

The PFAM PLDc_2 domain (PF13091) consensus sequence was used as input for iterative searches against the *Ae. aegypti* proteome using JackHMMR (https://www.ebi.ac.uk/Tools/hmmer/search/jackhmmer) to identify all *Ae. aegypti* PLDc_2 domain containing proteins. To generate a neighbor-joining tree, the PLDc_2 domain sequences from the identified genes and the Zuc orthologs in *Drosophila*, silkworm and mouse were aligned using M-Coffee (http://tcoffee.crg.cat/apps/tcoffee/do:mcoffee). One of the identified genes, AAEL022490, contains two PLDc_2 domains, both of which were used as separate inputs in the subsequent analysis. The same approach using the DEDDy type 5’-3’ exonuclease (cd06141) CDD-consensus sequence was used for the identification of *Ae. aegypti* Nbr.

For the alignment of full-length Zuc sequences, the entire sequence of *Ae. aegypti* Zuc as annotated on VectorBase (www.vectorbase.org), the Zuc sequence as determined by us from Aag2 cells and the sequences of Zuc orthologs in *Drosophila,* silkworm, mouse were aligned using M-Coffee with default settings and visualized using Jalview 2.11.0. Conservation of physicochemical properties at each position was analyzed in Jalview 2.11.0., and is based on the Analysis of Multiply Aligned Sequences (AMAS) method described in (8).

### RNA isolation

Cells were lysed in 1 mL RNA-Solv reagent (Omega Bio-tek), followed by RNA extraction through phase separation and isopropanol precipitation. RNA integrity was evaluated on EtBr stained agarose gels, and RNA concentration was determined using the Nanodrop ND-1000.

### RT-qPCR

For RT-qPCR analyses, DNaseI (Ambion)-treated RNA was reverse transcribed using the Taqman reverse transcriptase kit (Life Technologies) and PCR amplified in the presence of SYBR green, using the GoTaq qPCR system (Promega) according to the manufacturers’ recommendations. In knockdown experiments, expression levels of target genes were normalized to the expression of the housekeeping gene lysosomal aspartic protease (LAP) for *Ae. aegypti* samples or Ribosomal Protein L5 (RpL5) for *Ae. albopictus* samples and fold changes were calculated using the 2^-ΔΔCT^ method (9). Viral RNA levels were calculated using standard curves generated from a tenfold serial dilution series of a plasmid containing the SINV genome sequence. The standard curves were: log10 viral RNA level=-0.2041×CT + 10.572 (R^2^=0.9983) for Capsid and log10 viral RNA levels=-0.3029×CT + 13.027 (R^2^=0.9625) for Nsp4. The following primers were used:

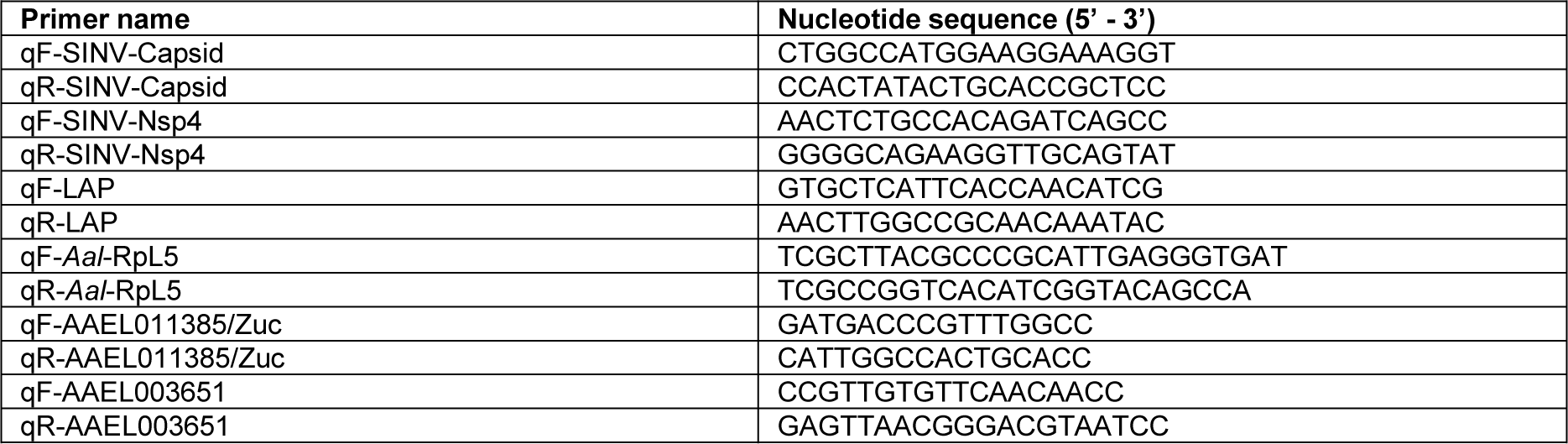
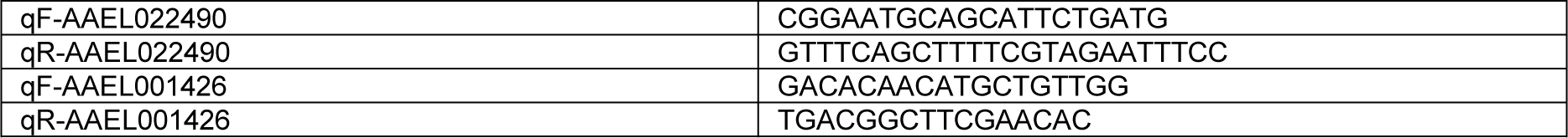

### Small RNA northern blotting

For small RNA northern blotting, 4-12 μg of RNA was size separated by denaturing urea polyacrylamide (15%) gel electrophoresis, transferred to nylon membranes and crosslinked using 1-ethyl-3-(3-dimethylaminopropyl)carbodiimide hydrochloride (EDC) crosslinking as described in (10). ^32^P labelled DNA probes (sequences shown below) were used to detect small RNAs.

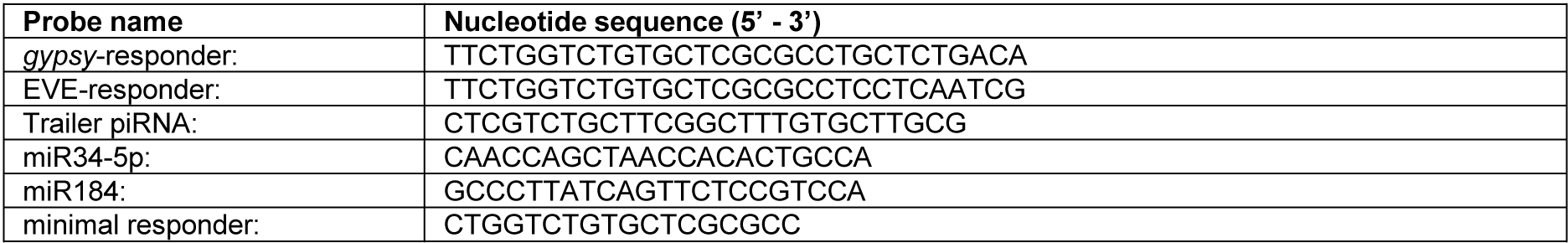

To generate the line graphs shown in Supplementary Figure S5C and S6C, northern blot signals were quantified at the center of each lane, using FIJI (11). Afterwards, peaks were manually aligned to correct for minor variations in size separation between samples. The average of five neighboring pixels was plotted to smoothen the line graph.

For the quantification of responder piRNA signals in Figures 3C-D, brightness and contrast was adjusted using the window/level function in FIJI to correct for background signal. Subsequently, the piRNA size range (24-33nt) was extrapolated from the EtBr stained marker, responder piRNA signal inside this range was quantified and normalized to the EtBr signal of rRNA to correct for variations in the amount of total RNA that was loaded on the gel.

### Generation of small RNA deep sequencing libraries

Small RNA deep sequencing libraries were generated using the NEBNext Small RNA Library Prep Set for Illumina (E7560, New England Biolabs), using 1 µg RNA as input. As piRNAs generally have 2’-*O*-methylated 3’ ends, we performed the 3’ adapter ligation for 18 hrs at 16°C to enhance the ligation efficiency of small RNAs bearing such modifications. The rest of the library preparation was performed in accordance with the manufacturer’s instructions. Libraries were sequenced on an Illumina Hiseq4000 machine by the GenomEast Platform (Strasbourg, France). Sequence data have been deposited in the NCBI sequence read archive under SRA accession SRP272125.

### Bioinformatic analyses of small RNA sequences

#### Quality control and small RNA mapping

RTA 2.7.3 and bcl2fastq were used for image analysis and base calling, and initial quality control was performed using FastQC. All subsequent manipulations were performed in Galaxy (12). First, the FASTX Clip adapter sequences tool (Galaxy version 1.0.3; default settings) was used to clip 3’ adapters from the small RNA sequence reads. Subsequently, reads were mapped to the corresponding recombinant Sindbis virus genomes, transposable element sequences extracted from TEFAM (originally downloaded from <u>tefam.biochem.vt.edu</u> on 10- 12-2019, now deposited at <u>github.com/RebeccaHalbach/Halbach_tapiR_2020/blob/master/Data/TEfam.fa</u>), *Ae. aegypti* transcripts (AaegL5.2 gene set, downloaded from VectorBase) and the Phasi Charoen like virus (PCLV) genome as sequenced from Aag2 cells (Genbank accession numbers KU936055, KU936056 and KU936057) (13). Reads were mapped using Bowtie allowing 1 mismatch, except for the mapping to interval and desert viruses and for 3’ end sharpness, downstream nucleotide bias, and phasing analyses, in which cases no mismatches were allowed. Small RNA libraries were normalized to the number of reads mapping to published pre-miRNA sequences deposited in miRBase v.21 without allowing mismatches.

#### Determining piRNA frequencies, size distributions and genome profiles

Size profiles were obtained by counting piRNA lengths after mapping, with the exception of the size profile of small RNAs obtained after PIWI IP (Figure S3), for which raw reads prior to mapping were analyzed from our previously published data (3). For genome distribution plots, the number of piRNA 5’ ends was determined for each position of the sequence to which small RNAs have been mapped.

For the analyses of piRNA production from the area of the non-coding reporter RNA downstream of the initiator/responder site versus the remainder of the viral genome shown in Figure 6G and Figure S7E, the SINV genome was defined as the area encompassing the 5’ UTR and the non-structural and structural polyproteins up to the start of the duplicated subgenomic promoter (nt 1-11385), and the trailer cassette as nt 11610-11753.

#### Size distribution and nucleotide biases for individual piRNAs

To generate heat maps of the length profile of piRNAs with a shared 5’ end published small RNA sequencing data was re-analyzed (2). SAM files were converted into interval files and piRNA-sized reads (25-30 nt) were filtered and separated according to the strand they mapped to. For each piRNA species, defined by a shared 5’ end, the percentage of reads with different lengths relative to the total read count was determined. piRNAs that were supported by less than 20 reads in total were discarded from the analysis. Subsequently, the percentages of read lengths per individual piRNA was imported into Multiple experiment viewer (v 4.9.0; tm4) and k-means clustering based on Pearson correlation was performed using the indicated number of clusters and a maximum number of iterations of 500. The ‘Construct hierarchical tree’ option was enabled. The graphical outputs of the clustering analyses were combined in Adobe Illustrator.

To analyze nucleotide biases within and downstream of piRNAs, only piRNA sequences that were supported by at least 20 reads and had a dominant piRNA length (at least 75% of all mapped reads have the same length) were considered. Combining the Get flanks (v1.0.0) and Get genomic DNA (v3.0.3) tools, the genomic sequence 5 nt downstream of that dominant piRNA length was extracted. Subsequently, each piRNA species was collapsed to one unique FastA sequence and its corresponding downstream sequence. The sequence logo was generated from these sequences using the Sequence logo generator (Galaxy version 3.5.0) The piRNA logos were cropped in Adobe Illustrator to show nucleotide biases of piRNA positions 1 to 5, 8 to 12, and the downstream positions +1 to +5.

#### Analysis of general piRNA sharpness

To analyze the effect of Zuc knockdown on vpiRNA 3’ ends, small RNAs mapping to the SINV genome up to the subgenomic promotor (nt 1-11385) were considered. For each of the six datasets (3x dsLuc and 3x dsZuc) available for the *gypsy*-25U virus, piRNA species were extracted by selecting only those piRNA start sites that were supported by at least fifty mature (25-30 nt) piRNA reads in the combined six datasets. For these piRNA positions, all reads in the size range of 25-38 nt were extracted from the original mapped data. As for the heat map analysis, the percentage of piRNA lengths was calculated for each piRNA species, defined by a shared 5’ end. To reduce noise due to low read count, only the 275 most abundant vpiRNA species were considered, which corresponded to approximately 160,000 reads in the combined six datasets. From the length distribution, a sharpness score was calculated for each piRNA species based on the maximum entropy (Smax = all reads have the same length) minus the observed Shannon entropy (Sobs) an approach similar to determining nucleotide biases (14): S_sharp_ = S_max_-S_obs_ = log2(14) - ∑^38^_n=25_ *p*_*n*_ * log _2_ *p*_*n*_

In this formula, the maximum score is defined as log2 of the number of different lengths that a piRNA could possibly have (25 to 38 nt); *p*_*n*_ is defined as the proportion of piRNA reads of a specific length (n) compared to all reads. The mean sharpness score for each piRNA was determined for the three control and three Zuc knockdown datasets separately and the 275 piRNAs were then ranked based on the mean sharpness score in the control knockdown from high score (sharp 3’ end) to lower scores (more diffuse 3’ end). Within this ranking an average of the mean sharpness scores was then calculated for 11 bins of 25 piRNA species each, both for the control and the Zuc knockdown. Finally, the difference in sharpness score (ΔSsharp) between control and Zuc knockdown was determined for each bin. The entire analysis was repeated for the six datasets of the *gypsy*-28U and *gypsy*-30U viruses and, since the sequence of the three viruses is identical up to the reporter cassette, the data was combined by calculating the mean + SEM of the ΔSsharp per bin obtained for each type of reporter virus. To determine statistical significance, the ΔSsharp was tested against the null hypothesis ΔSsharp=0. A two-sided student’s t-test was performed per bin and the hypothesis was rejected at *P* < 0.05. The sharpness score analysis for transposon mapping piRNAs was performed with the 4400 most expressed piRNAs (400 piRNAs per bin).

#### Analysis of piRNA phasing

piRNA phasing was analyzed for all vpiRNAs from the combined nine dsLuc datasets from the Zuc knockdown experiment (presented in Figure 4). For transposon derived piRNAs this analysis was performed on the combined three dsLuc datasets from the *gypsy-25U* viruses only to ensure that the number of transposon piRNAs reads analyzed was in the same order of magnitude as vpiRNAs. Only piRNAs that mapped to sequences common to all datasets were considered, piRNAs that map to the specific reporter sequences were excluded. First, only piRNA species, defined by a shared 5’ end, that were supported by at least 5 reads were selected. For these piRNAs, the distance of every individual piRNA 5’ end to up to 200 unique downstream piRNA 5’ ends, irrespective of read count, was determined using bedtools Closest bed (ClosestBed, Galaxy Version 2.30.0; settings: input A: piRNA 5’ ends of all piRNA reads; input B: piRNA 5’ ends of unique piRNA 5’ ends; report all ties (-b); only overlaps occurring on the **same** strand; report distance enabled; report upstream distances disabled (-d); report 200 closest hits (-k); ignore features in dataset B that overlap A disabled). The frequency of 5’- 5’ end distances was counted and plotted for a window of 20-150 nt (scatterplot in Figures 6A- B). The smoothened curves were determined using local regression (Loess; 10% of points fit; Kernel: Epanechnikov) in IBM SPSS v25.

### Generation of Zuc expression plasmids

The sequence encoding the Zuc protein was amplified from Aag2 cDNA using CloneAmp HiFi PCR Premix (Takara) and cloned into the pAWG vector (The Drosophila Gateway Vector Collection, Carnegie Science) using In-fusion technology (Takara). The following primers were used to amplify the vector and insert:

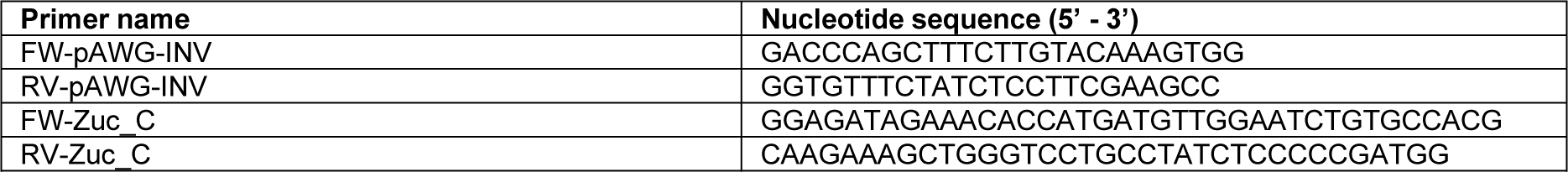

Subsequently, pAW3F-Zuc was generated by inverse PCR of pAWG-Zuc, which introduced the 3×flag tag to replace the eGFP-tag, followed by In-fusion cloning (Takara Biotech). The primers used for this inverse PCR are shown below (underlined sequence makes up the 3×flag tag, double underlined sequence is the 15 nt overlap required for In-fusion cloning):

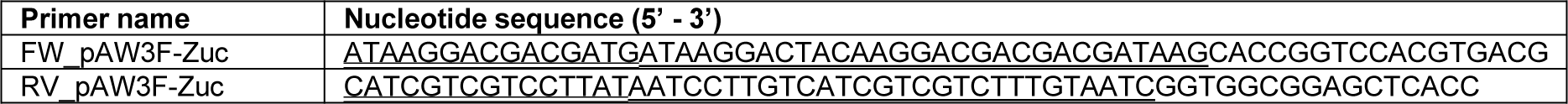

### Immunofluorescence

Approximately 5×10^5^ Aag2 cells were transfected with 1 µg of the pAW3F-Zuc expression vector using 1 µL X-tremeGENE HP DNA Transfection Reagent (Roche) according to the manufacturer’s instructions. 48 hrs later, cells were fixed on coverslips using 4% paraformaldehyde for 10 minutes at room temperature. Fixed cells were permeabilized in PBS-Triton (0.25%) for 10 minutes and blocked in 10% normal goat serum/0.3M Glycine in PBS-Tween (0.1%). Cells were stained with mouse anti-flag antibody (1:200, Sigma, F1804, RRID: AB_262044) for 1 hr at room temperature, followed by goat anti-mouse IgG Alexa fluor 568 (1:200, Invitrogen, A-11004, RRID: AB_2534072) for 1 hr at room temperature. Following antibody staining, mitochondria were stained using 200 µM Mitoview Green (Biotium), according to the manufacturer’s instruction. Lastly, Hoechst reagent was used to stain the nuclei and the coverslips were mounted onto microscope slides for imaging using the Zeiss LSM900 confocal microscope. In between all steps during the staining procedure, cells were washed three times with PBS.

### Co-immunoprecipitation and western blot

For immunoprecipitation of Zuc, ∼4.5 × 10^6^ Aag2 cells were seeded and after ∼16 hrs, 15 µg of the pAW3F-Zuc was transfected using X-tremeGENE HP DNA Transfection Reagent (Roche) according to the manufacturer’s instructions. Cells were lysed 48 hrs after transfection using 300 µL lysis buffer (10 mM Tris-HCl pH 7.5, 150 mM NaCl, 0.5 mM EDTA, 0.5% Igepal CA-630 [Sigma], 10% Glycerol, 1x cOmplete Protease Inhibitor [Roche], 1 mM PMSF). After incubation for 1 hr at 4°C with end-over-end rotation, lysates were centrifuged for 30 min at 15000 × g, 4°C. The supernatant was snap-frozen in liquid nitrogen and stored at -80°C for later use.

For immunoprecipitation, 15 µL M2-Flag bead slurry (Sigma) was equilibrated in lysis buffer, and, along with 450 µL dilution buffer (10 mM Tris-HCl pH 7.5, 150 mM NaCl, 0.5 mM EDTA, 1x cOmplete Protease Inhibitor [Roche], 1 mM PMSF), added to the thawed lysate. After incubation for 2 hrs at 4°C with end-over-end rotation, beads were washed thrice using 500 µL dilution buffer, before harvesting immunoprecipitates by boiling for 10 min in 20 µL 2× sample buffer (120 mM Tris/Cl pH 6.8, 20% glycerol, 4% SDS, 0.04% bromophenol blue, 10% β- mercaptoethanol). After boiling, samples were diluted by adding 20 µL lysis buffer.

Samples were resolved on 10% polyacrylamide gels and blotted to nitrocellulose membranes. The following antibodies, generated in our laboratory, were used for western blotting: rabbit-anti-Ago3, -Piwi4, -Piwi5 and -Piwi6 (all at 1:500) (3, 4). Additionally, mouse anti-flag (1:1000, Sigma, F1804, RRID: AB_262044) was used. Secondary antibodies were goat-anti-rabbit- IRdye800 [Li-cor; 926-32211, RRID: AB_621843] and goat-anti-mouse-IRdye680 [926-68070, RRID: AB_10956588].

## REFERENCES

1. Franz, A.W., Kantor, A.M., Passarelli, A.L. and Clem, R.J. (2015) Tissue Barriers to Arbovirus Infection in Mosquitoes. Viruses, 7, 3741–3767.

2. Bronkhorst, A.W. and van Rij, R.P. (2014) The long and short of antiviral defense: small RNA-based immunity in insects. Curr Opin Virol, 7, 19–28.

3. Miesen, P., Joosten, J. and van Rij, R.P. (2016) PIWIs Go Viral: Arbovirus-Derived piRNAs in Vector Mosquitoes. PLoS Pathog, 12, e1006017.

4. Ozata, D.M., Gainetdinov, I., Zoch, A., O’Carroll, D. and Zamore, P.D. (2019) PIWI-interacting RNAs: small RNAs with big functions. Nat Rev Genet, 20, 89–108.

5. Czech, B. and Hannon, G.J. (2016) One Loop to Rule Them All: The Ping-Pong Cycle and piRNA-Guided Silencing. Trends Biochem Sci, 41, 324–337.

6. Gainetdinov, I., Colpan, C., Arif, A., Cecchini, K. and Zamore, P.D. (2018) A Single Mechanism of Biogenesis, Initiated and Directed by PIWI Proteins, Explains piRNA Production in Most Animals. Mol Cell, 71, 775–790 e775.

7. Han, B.W., Wang, W., Li, C.J., Weng, Z.P. and Zamore, P.D. (2015) piRNA-guided transposon cleavage initiates Zucchini- dependent, phased piRNA production. Science, 348, 817–821.

8. Mohn, F., Handler, D. and Brennecke, J. (2015) piRNA-guided slicing specifies transcripts for Zucchini-dependent, phased piRNA biogenesis. Science, 348, 812–817.

9. Horwich, M.D., Li, C., Matranga, C., Vagin, V., Farley, G., Wang, P. and Zamore, P.D. (2007) The Drosophila RNA methyltransferase, DmHen1, modifies germline piRNAs and single-stranded siRNAs in RISC. Curr Biol, 17, 1265–1272.

10. Saito, K., Sakaguchi, Y., Suzuki, T., Suzuki, T., Siomi, H. and Siomi, M.C. (2007) Pimet, the Drosophila homolog of HEN1, mediates 2’-O-methylation of Piwi- interacting RNAs at their 3’ ends. Genes Dev, 21, 1603–1608.

11. Hayashi, R., Schnabl, J., Handler, D., Mohn, F., Ameres, S.L. and Brennecke, J. (2016) Genetic and mechanistic diversity of piRNA 3’-end formation. Nature, 539, 588–592.

12. Feltzin, V.L., Khaladkar, M., Abe, M., Parisi, M., Hendriks, G.J., Kim, J. and Bonini, N.M. (2015) The exonuclease Nibbler regulates age-associated traits and modulates piRNA length in Drosophila. Aging Cell, 14, 443–452.

13. Wang, H., Ma, Z., Niu, K., Xiao, Y., Wu, X., Pan, C., Zhao, Y., Wang, K., Zhang, Y. and Liu, N. (2016) Antagonistic roles of Nibbler and Hen1 in modulating piRNA 3’ ends in Drosophila. Development, 143, 530–539.

14. Sienski, G., Donertas, D. and Brennecke, J. (2012) Transcriptional silencing of transposons by Piwi and maelstrom and its impact on chromatin state and gene expression. Cell, 151, 964–980.

15. Le Thomas, A., Rogers, A.K., Webster, A., Marinov, G.K., Liao, S.E., Perkins, E.M., Hur, J.K., Aravin, A.A. and Toth, K.F. (2013) Piwi induces piRNA-guided transcriptional silencing and establishment of a repressive chromatin state. Genes Dev, 27, 390–399.

16. Gunawardane, L.S., Saito, K., Nishida, K.M., Miyoshi, K., Kawamura, Y., Nagami, T., Siomi, H. and Siomi, M.C. (2007) A slicer- mediated mechanism for repeat-associated siRNA 5’ end formation in Drosophila. Science, 315, 1587–1590.

17. Brennecke, J., Aravin, A.A., Stark, A., Dus, M., Kellis, M., Sachidanandam, R. and Hannon, G.J. (2007) Discrete small RNA- generating loci as master regulators of transposon activity in Drosophila. Cell, 128, 1089–1103.

18. Malone, C.D., Brennecke, J., Dus, M., Stark, A., McCombie, W.R., Sachidanandam, R. and Hannon, G.J. (2009) Specialized piRNA pathways act in germline and somatic tissues of the Drosophila ovary. Cell, 137, 522–535.

19. Li, C., Vagin, V.V., Lee, S., Xu, J., Ma, S., Xi, H., Seitz, H., Horwich, M.D., Syrzycka, M., Honda, B.M. et al. (2009) Collapse of germline piRNAs in the absence of Argonaute3 reveals somatic piRNAs in flies. Cell, 137, 509–521.

20. Lewis, S.H., Quarles, K.A., Yang, Y., Tanguy, M., Frezal, L., Smith, S.A., Sharma, P.P., Cordaux, R., Gilbert, C., Giraud, I. et al. (2018) Pan-arthropod analysis reveals somatic piRNAs as an ancestral defence against transposable elements. Nat Ecol Evol, 2, 174–181.

21. Akbari, O.S., Antoshechkin, I., Amrhein, H., Williams, B., Diloreto, R., Sandler, J. and Hay, B.A. (2013) The developmental transcriptome of the mosquito Aedes aegypti, an invasive species and major arbovirus vector. G3 (Bethesda), 3, 1493–1509.

22. Campbell, C.L., Black, W.C.t., Hess, A.M. and Foy, B.D. (2008) Comparative genomics of small RNA regulatory pathway components in vector mosquitoes. BMC Genomics, 9, 425.

23. Lewis, S.H., Salmela, H. and Obbard, D.J. (2016) Duplication and Diversification of Dipteran Argonaute Genes, and the Evolutionary Divergence of Piwi and Aubergine. Genome Biol Evol, 8, 507–518.

24. Miesen, P., Girardi, E. and van Rij, R.P. (2015) Distinct sets of PIWI proteins produce arbovirus and transposon-derived piRNAs in Aedes aegypti mosquito cells. Nucleic Acids Res, 43, 6545–6556.

25. Joosten, J., Miesen, P., Taskopru, E., Pennings, B., Jansen, P., Huynen, M.A., Vermeulen, M. and Van Rij, R.P. (2019) The Tudor protein Veneno assembles the ping-pong amplification complex that produces viral piRNAs in Aedes mosquitoes. Nucleic Acids Res, 47, 2546–2559.

26. Morazzani, E.M., Wiley, M.R., Murreddu, M.G., Adelman, Z.N. and Myles, K.M. (2012) Production of virus-derived ping-pong- dependent piRNA-like small RNAs in the mosquito soma. PLoS Pathog, 8, e1002470.

27. Vodovar, N., Bronkhorst, A.W., van Cleef, K.W., Miesen, P., Blanc, H., van Rij, R.P. and Saleh, M.C. (2012) Arbovirus-derived piRNAs exhibit a ping-pong signature in mosquito cells. PLoS One, 7, e30861.

28. Palatini, U., Miesen, P., Carballar-Lejarazu, R., Ometto, L., Rizzo, E., Tu, Z., van Rij, R.P. and Bonizzoni, M. (2017) Comparative genomics shows that viral integrations are abundant and express piRNAs in the arboviral vectors Aedes aegypti and Aedes albopictus. BMC Genomics, 18, 512.

29. Whitfield, Z.J., Dolan, P.T., Kunitomi, M., Tassetto, M., Seetin, M.G., Oh, S., Heiner, C., Paxinos, E. and Andino, R. (2017) The Diversity, Structure, and Function of Heritable Adaptive Immunity Sequences in the Aedes aegypti Genome. Curr Biol, 27, 3511–3519 e3517.

30. Suzuki, Y., Frangeul, L., Dickson, L.B., Blanc, H., Verdier, Y., Vinh, J., Lambrechts, L. and Saleh, M.C. (2017) Uncovering the Repertoire of Endogenous Flaviviral Elements in Aedes Mosquito Genomes. J Virol, 91.

31. Aguiar, E., de Almeida, J.P.P., Queiroz, L.R., Oliveira, L.S., Olmo, R.P., de Faria, I., Imler, J.L., Gruber, A., Matthews, B.J. and Marques, J.T. (2020) A single unidirectional piRNA cluster similar to the flamenco locus is the major source of EVE-derived transcription and small RNAs in Aedes aegypti mosquitoes. RNA, 26, 581–594.

32. Crava, C.M., Varghese, F.S., Pischedda, E., Halbach, R., Palatini, U., Marconcini, M., Gasmi, L., Redmond, S., Afrane, Y., Ayala, D. et al. (2021) Population genomics in the arboviral vector Aedes aegypti reveals the genomic architecture and evolution of endogenous viral elements. Mol Ecol, 30, 1594–1611.

33. Ter Horst, A.M., Nigg, J.C., Dekker, F.M. and Falk, B.W. (2019) Endogenous Viral Elements Are Widespread in Arthropod Genomes and Commonly Give Rise to PIWI-Interacting RNAs. J Virol, 93.

34. Tassetto, M., Kunitomi, M., Whitfield, Z.J., Dolan, P.T., Sanchez-Vargas, I., Garcia-Knight, M., Ribiero, I., Chen, T., Olson, K.E. and Andino, R. (2019) Control of RNA viruses in mosquito cells through the acquisition of vDNA and endogenous viral elements. Elife, 8.

35. Suzuki, Y., Baidaliuk, A., Miesen, P., Frangeul, L., Crist, A.B., Merkling, S.H., Fontaine, A., Lequime, S., Moltini-Conclois, I., Blanc, H. et al. (2020) Non-retroviral Endogenous Viral Element Limits Cognate Virus Replication in Aedes aegypti Ovaries. Curr Biol, 30, 3495–3506 e3496.

36. Blankenberg, D., Gordon, A., Von Kuster, G., Coraor, N., Taylor, J., Nekrutenko, A. and Galaxy, T. (2010) Manipulation of FASTQ data with Galaxy. Bioinformatics, 26, 1783–1785.

37. Langmead, B. and Salzberg, S.L. (2012) Fast gapped-read alignment with Bowtie 2. Nat Methods, 9, 357–359.

38. Halbach, R., Miesen, P., Joosten, J., Taskopru, E., Rondeel, I., Pennings, B., Vogels, C.B.F., Merkling, S.H., Koenraadt, C.J., Lambrechts, L. et al. (2020) A satellite repeat-derived piRNA controls embryonic development of Aedes. Nature, 580, 274–277.

39. Ipsaro, J.J., Haase, A.D., Knott, S.R., Joshua-Tor, L. and Hannon, G.J. (2012) The structural biochemistry of Zucchini implicates it as a nuclease in piRNA biogenesis. Nature, 491, 279–283.

40. Nishimasu, H., Ishizu, H., Saito, K., Fukuhara, S., Kamatani, M.K., Bonnefond, L., Matsumoto, N., Nishizawa, T., Nakanaga, K., Aoki, J. et al. (2012) Structure and function of Zucchini endoribonuclease in piRNA biogenesis. Nature, 491, 284–287.

41. Izumi, N., Shoji, K., Suzuki, Y., Katsuma, S. and Tomari, Y. (2020) Zucchini consensus motifs determine the mechanism of pre- piRNA production. Nature, 578, 311–316.

42. Strauss, J.H. and Strauss, E.G. (1994) The alphaviruses: gene expression, replication, and evolution. Microbiol Rev, 58, 491–562.

43. Fredericks, A.C., Russell, T.A., Wallace, L.E., Davidson, A.D., Fernandez-Sesma, A. and Maringer, K. (2019) Aedes aegypti (Aag2)-derived clonal mosquito cell lines reveal the effects of pre-existing persistent infection with the insect-specific bunyavirus Phasi Charoen-like virus on arbovirus replication. PLoS Negl Trop Dis, 13, e0007346.

44. Joosten, J., Taskopru, E., Jansen, P., Pennings, B., Vermeulen, M. and Van Rij, R.P. (2021) PIWI proteomics identifies Atari and Pasilla as piRNA biogenesis factors in Aedes mosquitoes. Cell Rep, 35, 109073.

45. Girardi, E., Miesen, P., Pennings, B., Frangeul, L., Saleh, M.C. and van Rij, R.P. (2017) Histone-derived piRNA biogenesis depends on the ping-pong partners Piwi5 and Ago3 in Aedes aegypti. Nucleic Acids Res, 45, 4881–4892.

46. Schnettler, E., Donald, C.L., Human, S., Watson, M., Siu, R.W.C., McFarlane, M., Fazakerley, J.K., Kohl, A. and Fragkoudis, R. (2013) Knockdown of piRNA pathway proteins results in enhanced Semliki Forest virus production in mosquito cells. J Gen Virol, 94, 1680–1689.

47. Reuter, M., Berninger, P., Chuma, S., Shah, H., Hosokawa, M., Funaya, C., Antony, C., Sachidanandam, R. and Pillai, R.S. (2011) Miwi catalysis is required for piRNA amplification-independent LINE1 transposon silencing. Nature, 480, 264–267.

48. Betting, V., Joosten, J., Halbach, R., Thaler, M., Miesen, P. and Van Rij, R.P. (2020) A regulatory network of a piRNA and lncRNA initiates responder and trailer piRNA formation during embryonic development of Aedes mosquitoes. bioRxiv, 2020.2003.2023.003038.

49. Pane, A., Wehr, K. and Schupbach, T. (2007) zucchini and squash encode two putative nucleases required for rasiRNA production in the Drosophila germline. Developmental cell, 12, 851–862.

50. Huang, H., Li, Y., Szulwach, K.E., Zhang, G., Jin, P. and Chen, D. (2014) AGO3 Slicer activity regulates mitochondria-nuage localization of Armitage and piRNA amplification. J Cell Biol, 206, 217–230.

51. Haase, A.D., Fenoglio, S., Muerdter, F., Guzzardo, P.M., Czech, B., Pappin, D.J., Chen, C., Gordon, A. and Hannon, G.J. (2010) Probing the initiation and effector phases of the somatic piRNA pathway in Drosophila. Genes Dev, 24, 2499–2504.

52. Xie, W., Sowemimo, I., Hayashi, R., Wang, J., Burkard, T.R., Brennecke, J., Ameres, S.L. and Patel, D.J. (2020) Structure- function analysis of microRNA 3’-end trimming by Nibbler. Proc Natl Acad Sci U S A, 117, 30370–30379.

53. Han, B.W., Hung, J.H., Weng, Z., Zamore, P.D. and Ameres, S.L. (2011) The 3’-to-5’ exoribonuclease Nibbler shapes the 3’ ends of microRNAs bound to Drosophila Argonaute1. Current biology : CB, 21, 1878–1887.

54. Liu, N., Abe, M., Sabin, L.R., Hendriks, G.J., Naqvi, A.S., Yu, Z., Cherry, S. and Bonini, N.M. (2011) The exoribonuclease Nibbler controls 3’ end processing of microRNAs in Drosophila. Current biology : CB, 21, 1888–1893.

55. Etebari, K., Osei-Amo, S., Blomberg, S.P. and Asgari, S. (2015) Dengue virus infection alters post-transcriptional modification of microRNAs in the mosquito vector Aedes aegypti. Sci Rep, 5, 15968.

56. Li, S., Mead, E.A., Liang, S. and Tu, Z. (2009) Direct sequencing and expression analysis of a large number of miRNAs in Aedes aegypti and a multi-species survey of novel mosquito miRNAs. BMC Genomics, 10, 581.

57. Izumi, N., Shoji, K., Sakaguchi, Y., Honda, S., Kirino, Y., Suzuki, T., Katsuma, S. and Tomari, Y. (2016) Identification and Functional Analysis of the Pre-piRNA 3’ Trimmer in Silkworms. Cell, 164, 962–973.

58. Tang, W., Tu, S., Lee, H.C., Weng, Z. and Mello, C.C. (2016) The RNase PARN-1 Trims piRNA 3’ Ends to Promote Transcriptome Surveillance in C. elegans. Cell, 164, 974–984.

59. Palatini, U., Miesen, P., Carballar-Lejarazu, R., Ometto, L., Rizzo, E., Tu, Z., van Rij, R.P. and Bonizzoni, M. (2017) Comparative genomics shows that viral integrations are abundant and express piRNAs in the arboviral vectors Aedes aegypti and Aedes albopictus. BMC Genomics, 18, 512.

60. Miesen, P., Girardi, E. and van Rij, R.P. (2015) Distinct sets of PIWI proteins produce arbovirus and transposon-derived piRNAs in Aedes aegypti mosquito cells. Nucleic Acids Res, 43, 6545–6556.

61. Joosten, J., Miesen, P., Taskopru, E., Pennings, B., Jansen, P., Huynen, M.A., Vermeulen, M. and Van Rij, R.P. (2019) The Tudor protein Veneno assembles the ping-pong amplification complex that produces viral piRNAs in Aedes mosquitoes. Nucleic Acids Res, 47, 2546–2559.

62. Halbach, R., Miesen, P., Joosten, J., Taskopru, E., Rondeel, I., Pennings, B., Vogels, C.B.F., Merkling, S.H., Koenraadt, C.J., Lambrechts, L. et al. (2020) A satellite repeat-derived piRNA controls embryonic development of Aedes. Nature, 580, 274–277.

63. Hahn, C.S., Hahn, Y.S., Braciale, T.J. and Rice, C.M. (1992) Infectious Sindbis virus transient expression vectors for studying antigen processing and presentation. Proc Natl Acad Sci U S A, 89, 2679–2683.

64. Saleh, M.C., Tassetto, M., van Rij, R.P., Goic, B., Gausson, V., Berry, B., Jacquier, C., Antoniewski, C. and Andino, R. (2009) Antiviral immunity in Drosophila requires systemic RNA interference spread. Nature, 458, 346–350.

65. Vodovar, N., Bronkhorst, A.W., van Cleef, K.W., Miesen, P., Blanc, H., van Rij, R.P. and Saleh, M.C. (2012) Arbovirus-derived piRNAs exhibit a ping-pong signature in mosquito cells. PLoS One, 7, e30861.

66. Livingstone, C.D. and Barton, G.J. (1993) Protein sequence alignments: a strategy for the hierarchical analysis of residue conservation. Comput Appl Biosci, 9, 745–756.

67. Livak, K.J. and Schmittgen, T.D. (2001) Analysis of relative gene expression data using real-time quantitative PCR and the 2(-Delta Delta C(T)) Method. Methods, 25, 402–408.

68. Pall, G.S. and Hamilton, A.J. (2008) Improved northern blot method for enhanced detection of small RNA. Nat Protoc, 3, 1077–1084.

69. Rueden, C.T., Schindelin, J., Hiner, M.C., DeZonia, B.E., Walter, A.E., Arena, E.T. and Eliceiri, K.W. (2017) ImageJ2: ImageJ for the next generation of scientific image data. BMC Bioinformatics, 18, 529.

70. Blankenberg, D., Gordon, A., Von Kuster, G., Coraor, N., Taylor, J., Nekrutenko, A. and Galaxy, T. (2010) Manipulation of FASTQ data with Galaxy. Bioinformatics, 26, 1783–1785.

71. Maringer, K., Yousuf, A., Heesom, K.J., Fan, J., Lee, D., Fernandez-Sesma, A., Bessant, C., Matthews, D.A. and Davidson, A.D. (2017) Proteomics informed by transcriptomics for characterising active transposable elements and genome annotation in Aedes aegypti. Bmc Genomics, 18, 101.

72. Crooks, G.E., Hon, G., Chandonia, J.M. and Brenner, S.E. (2004) WebLogo: a sequence logo generator. Genome Res, 14, 1188–1190.

## Supplemental References

1. Palatini, U., Miesen, P., Carballar-Lejarazu, R., Ometto, L., Rizzo, E., Tu, Z., van Rij, R.P. and Bonizzoni, M. (2017) Comparative genomics shows that viral integrations are abundant and express piRNAs in the arboviral vectors Aedes aegypti and Aedes albopictus. BMC Genomics, 18, 512.

2. Miesen, P., Girardi, E. and van Rij, R.P. (2015) Distinct sets of PIWI proteins produce arbovirus and transposon-derived piRNAs in Aedes aegypti mosquito cells. Nucleic Acids Res, 43, 6545–6556.

3. Joosten, J., Miesen, P., Taskopru, E., Pennings, B., Jansen, P., Huynen, M.A., Vermeulen, M. and Van Rij, R.P. (2019) The Tudor protein Veneno assembles the ping-pong amplification complex that produces viral piRNAs in Aedes mosquitoes. Nucleic Acids Res, 47, 2546–2559.

4. Halbach, R., Miesen, P., Joosten, J., Taskopru, E., Rondeel, I., Pennings, B., Vogels, C.B.F., Merkling, S.H., Koenraadt, C.J., Lambrechts, L. et al. (2020) A satellite repeat-derived piRNA controls embryonic development of Aedes. Nature, 580, 274–277.

5. Hahn, C.S., Hahn, Y.S., Braciale, T.J. and Rice, C.M. (1992) Infectious Sindbis virus transient expression vectors for studying antigen processing and presentation. Proc Natl Acad Sci U S A, 89, 2679–2683.

6. Saleh, M.C., Tassetto, M., van Rij, R.P., Goic, B., Gausson, V., Berry, B., Jacquier, C., Antoniewski, C. and Andino, R. (2009) Antiviral immunity in Drosophila requires systemic RNA interference spread. Nature, 458, 346–350.

7. Vodovar, N., Bronkhorst, A.W., van Cleef, K.W., Miesen, P., Blanc, H., van Rij, R.P. and Saleh, M.C. (2012) Arbovirus-derived piRNAs exhibit a ping-pong signature in mosquito cells. PLoS One, 7, e30861.

8. Livingstone, C.D. and Barton, G.J. (1993) Protein sequence alignments: a strategy for the hierarchical analysis of residue conservation. Comput Appl Biosci, 9, 745–756.

9. Livak, K.J. and Schmittgen, T.D. (2001) Analysis of relative gene expression data using real-time quantitative PCR and the 2(-Delta Delta C(T)) Method. Methods, 25, 402–408.

10. Pall, G.S. and Hamilton, A.J. (2008) Improved northern blot method for enhanced detection of small RNA. Nat Protoc, 3, 1077–1084.

11. Rueden, C.T., Schindelin, J., Hiner, M.C., DeZonia, B.E., Walter, A.E., Arena, E.T. and Eliceiri, K.W. (2017) ImageJ2: ImageJ for the next generation of scientific image data. BMC Bioinformatics, 18, 529.

12. Blankenberg, D., Gordon, A., Von Kuster, G., Coraor, N., Taylor, J., Nekrutenko, A. and Galaxy, T. (2010) Manipulation of FASTQ data with Galaxy. Bioinformatics, 26, 1783–1785.

13. Maringer, K., Yousuf, A., Heesom, K.J., Fan, J., Lee, D., Fernandez-Sesma, A., Bessant, C., Matthews, D.A. and Davidson, A.D. (2017) Proteomics informed by transcriptomics for characterising active transposable elements and genome annotation in Aedes aegypti. Bmc Genomics, 18, 101.

14. Crooks, G.E., Hon, G., Chandonia, J.M. and Brenner, S.E. (2004) WebLogo: a sequence logo generator. Genome Res, 14, 1188–1190.

